# The adaptive molecular landscape of reprogrammed telomeric sequences

**DOI:** 10.1101/2025.06.24.661269

**Authors:** Melania D’Angiolo, Benjamin P. Barré, Sakshi Khaiwal, Julia Muenzner, Johan Hallin, Matteo De Chiara, Nicolò Tellini, Jonas Warringer, Markus Ralser, Eric Gilson, Gianni Liti

## Abstract

Telomeric sequences vary across the tree of life and intimately co-evolve with telomere-binding protein complexes. However, the molecular mechanisms allowing organisms to adapt to new telomeric sequences are difficult to gauge from extant species. Here, we reprogrammed multiple yeast lines to human-like telomeric repeats to unveil their molecular and fitness response to novel telomeres. Initially, the exchange of telomere sequences resulted in genome instability, proteome remodelling and severe fitness decline. However, adaptive evolution experiments selected for repeated mutations that drove adaptation to the humanized telomeres. These consisted of the recurrent amplification of the telomere-binding protein *TBF1*, by complex aneuploidies, or in repeated mutations that attenuate the DNA damage response. Overall, our results outline a response that defines the adaptive molecular landscape to novel telomeric sequences.

## Introduction

The correct maintenance of the genome is of fundamental importance in order to prevent instability and degeneration. In most eukaryotes, the genome is organized in linear segments whose extremities are exposed to degradation and need to be protected. This function is performed by ribonucleoproteins called “telomeres”, consisting of a repeated DNA sequence and its associated proteins (Greider & Blackburn, 1987). Telomeres are maintained via the specialised enzyme telomerase. This enzyme is a reverse transcriptase that synthesizes new repeats using a RNA moiety as template (Chan & Blackburn, 2002; De Lange, 2005; Greider & Blackburn, 1989; Lundblad & Szostak, 1989; Marcand et al., 1997).

Despite variations in their DNA sequence having occurred during evolution, the function of telomeres in genome maintenance is remarkably conserved across the tree of life. In fact, all telomeric DNA motifs share common characteristics, like being short, tandemly-repeated and G-rich, and the principles of telomere biology are widely conserved even among distant taxa (Kupiec, 2014). Nevertheless, variations in both telomeric sequence and length occur (Lyčka et al., 2024; Whittemore et al., 2019). The vertebrate-like TTAGGG (T_2_AG_3_) repeat is the most widespread, being present in the majority of Metazoa, but also in some classes of protists, fungi and plants, and is thought to be the ancestral unit from which all the others evolved (Fulnečková et al., 2013). Variations of this motif are observed in other taxa, like TTTAGGG in plants (Peska & Garcia, 2020), TTAGG in insects (Kuznetsova et al., 2020), TTAGGC in nematodes (Zetka & Müller, 1996) and degenerate motifs in *Saccharomyces* yeasts (Teixeira & Gilson, 2005). In other organisms, telomeric repeat units can be very long and complex, like in species of the genus *Candida* or in fission yeasts (Sepsiova et al., 2016; Peska et al., 2021). Therefore, telomeric DNA sequences have changed at multiple times during evolution, but how organisms subsequently adapt to accomodate and capitalize on these changes remains uncharacterised.

Telomeric DNA varies in length with a difference of several orders of magnitude across species and extensive within-species variation. Telomere length ranges from 20 bp in the ciliate *Oxytricha trifallax* (Cavalcanti et al., 2004), 150-600 bp in budding yeasts (D’Angiolo et al., 2023; Garrido et al., 2025; O’Donnell et al., 2023), to 5-15 kb in humans (Karimian et al., 2024) and over 100 kb in some mice species (Hemann & Greider, 2000). The budding yeast *Saccharomyces cerevisiae* has been a leading model in telomere biology. *S. cerevisiae* telomeres are constituted by degenerate TG_1-3_ repeats (Wellinger & Zakian, 2012). Moreover, yeast telomeres are flanked by repetitive subtelomeric DNA like the X and Y′ elements, and interstitial telomeric sequences (ITS) that are often located between them, adding further complexity (Louis, 1995; Wellinger & Zakian, 2012). *S. cerevisiae* cells can survive for several cell divisions after the complete loss of telomerase, and often generate stable survivors that can replicate indefinitely thanks to recombination-based maintenance of telomeres (Lundblad & Blackburn, 1993).

It is possible to generate yeasts with human-like T_2_AG_3_ telomeres by engineering the telomerase RNA template encoded by the gene *TLC1* (Henning et al., 1998; Forstemann & Lingner, 2001). Such a humanization of yeast telomeres entails a switch in telomere-binding proteins, in which Rap1p, originally bound to native TG_1-3_ repeats, is replaced by Tbf1p, a regulatory factor known to bind TTAGGG repeats in the yeast genome (Brigati et al., 1993; Koering et al., 2000; Berthiau et al., 2006). Humanized telomeres are dysfunctional and show deregulation of their length and loss of subtelomeric transcriptional silencing. Furthermore, they trigger the activation of the DNA damage response (DDR) with consequent cell cycle arrest (Alexander & Zakian, 2003; Auriche et al., 2008; Bah et al., 2004; Bah et al., 2011; Brevet et al., 2003; Di Domenico et al., 2009, 2013; Fukunaga et al., 2012; Ribaud et al., 2012).

In this study, we investigated the evolutionary response of yeast cells to a sudden reprogramming of all telomeres by propagating them through experimental evolution. We sequentially combined two experimental evolution paradigms: first, we propagated cells over many mitotic generations through mutation accumulation lines (MAL) to minimize selection and study the long-term effects of telomere reprogramming. Next, we submitted MAL to adaptive evolution (AEL) by repeated serial transfers of large populations in which multiple clones are competing against each other, to map mutations that counteract the fitness decline. The latter procedure ensures that if some clones develop a beneficial mutation, they will outcompete the others and lead the mutation to fixation. During MAL, yeasts with humanized telomers showed a progressive fitness decay accompanied by genome instability and partial proteomic remodelling, but recovered near-wild-type fitness after AEL by amplifying *TBF1* or modulating the DNA damage response.

## Results

### An evolutionary framework of novel telomeres

Previous work has shown that *S. cerevisiae* yeasts engineered to carry point mutations in the *TLC1*-encoded 16 nt telomerase RNA template have growth and chromosome segregation defects (Lin et al., 2004), suggesting that this motif is under strong evolutionary constraints that limit its freedom to vary in nature. To quantify these constraints, we examined the template and its flanking regions in 8 *Saccharomyces* species spanning nearly 20 MY of evolution (Shen et al., 2018). Our multiple sequence alignment revealed that while flanking regions show variation that reflects the species evolutionary relationships the template region is extremely conserved, with only two variants: in the first and last nt positions (**Fig. 1a**). We further analyzed intraspecific variation in *TLC1* in a collection of 3380 natural *S. cerevisiae* isolates to explore recent mutations across the ecological and geographic range of this model species. We found two rare variants, again in the first or the last nt of the template, with minor allele frequencies of 0.002 and 0.0003 respectively (**Fig. 1a**). Thus, purifying selection acting on the *TLC1* sequence in natural yeast lineages is remarkably strong.

**Fig. 1.**
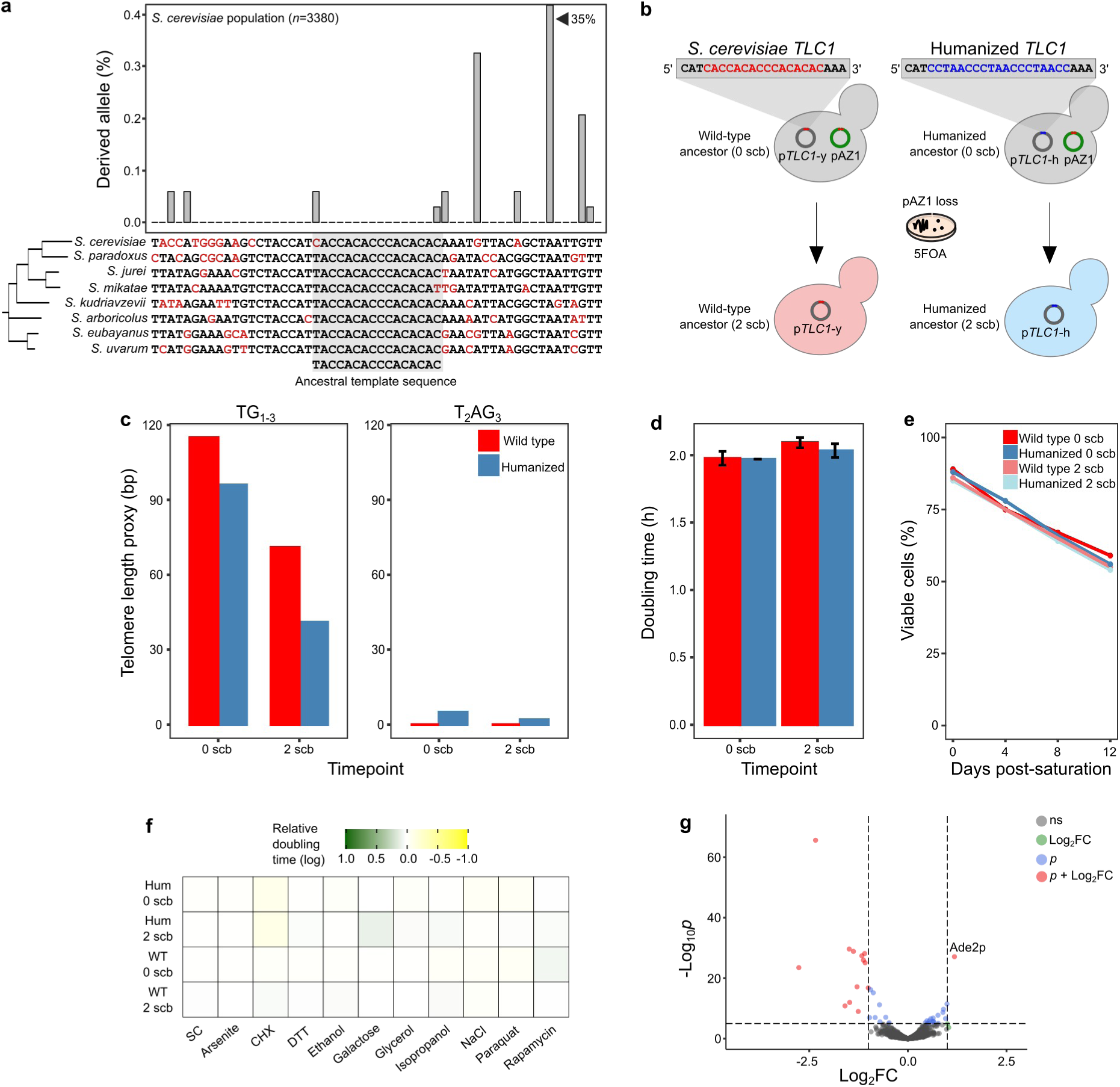
*TLC1* evolution and reprogramming. **a,** Natural sequence variation of the telomerase RNA template (*TLC1*). The barplot represents the percentage of *S. cerevisiae* strains carrying a derived allele at each position of the alignment (*n*=3380). The alignment is based on the 8 species of the *Saccharomyces* genus. Black letters denote ancestral alleles while red letters denote the inferred derived alleles. The upper panel represents an intra-specific comparison, while the bottom panel represents an inter-specific comparison. **b**, Schematics of the plasmids contained in the ancestor strains at 0 and 2 scb. **c-e**, Telomere length, population doubling time and survival rate of wild-type and humanized ancestors at 0 and 2 scb. Barplots: mean + standard deviation. Telomere length measurements were performed in one replicate. Population doubling time measurements were performed in triplicate. Survival rate measurements were performed in one replicate. **f,** Relative doubling time of wild-type and humanized ancestors at 0 and 2 scb. Colours indicate growth normalized using wild-type at 2 scb (MJD5). Relative doubling time measurements were performed in 24 replicates for SC and 12 replicates for all the other conditions. **g**, Volcano plot showing differentially expressed proteins in the humanized ancestor at 2 scb as compared to wild-type 2 scb. Proteins supported by fold-change (x), *p*-value (y) or both are indicated in color, but only the ones supported by both (red) were retained for analysis.

In order to reconfigure *S. cerevisiae* telomere repeats, we used a genetic system that generates isogenic strains with either wild-type or humanized telomeres (Bah et al., 2004; Bah et al., 2011) (**Table S1**). In brief, these two strains differ by a low-copy centromeric plasmid with either a native (p*TLC1*-y) or humanized (p*TLC1*-h) copy of *TLC1*, with the chromosomal *TLC1* having been deleted from its native locus. These strains contain an additional plasmid with a copy of *TLC1*-y (pAZ1) that serves as a back-up and pAZ1 loss can be controlled by plating cells in a selective medium (**Fig. 1b**).

We initiated the telomere reprogramming by streaking the ancestors on 5-FOA for two single-cell bottlenecks (scb) to select for *URA3*-containing pAZ1 loss. We investigated the effect of pAZ1 loss on telomere DNA length and composition, as well as on mitotic cell growth and lifespan. First, we sequenced the ancestors’ genomes before (0 scb) and after pAZ1 loss (2 scb) to estimate telomere length and composition from short-read data (D’Angiolo et al., 2023). The TG_1-3_ telomere length decreased upon pAZ1 loss, for both the wild-type and humanized ancestor, consistent with telomere length being affected by the copy number of *TLC1* (Zappulla & Cech, 2004). The wild-type did not contain T_2_AG_3_ repeats, while a small amount was detected (5% T_2_AG_3_ reads) in the humanized ancestor at both timepoints (**Fig. 1c and Table S2**). The rare T_2_AG_3_ telomeric repeats and the loss of pAZ1 did not immediately affect the cell’s doubling time across several conditions or chronological lifespan (**Fig. 1d-f, Tables S5-7**).

Next, we investigated the impact of pAZ1 loss on the global proteome, using liquid chromatography tandem mass spectrometry. We only detected 4 differentially expressed proteins in the humanized ancestor at 0 scb as compared to the wild-type (**Table S11**). However, we identified 14 differentially expressed proteins in the humanized ancestor at 2 scb. In humanized yeasts, *ADE2* is inactivated by a loss-of-function mutation in its native locus and a functional copy is inserted into the right subtelomere of chrV and acts as subtelomeric transcriptional silencing reporter. The replacement of Rap1p by Tbf1p results in transcriptional derepression of subtelomeric genes most likely due to the lack of the deacetylases Sir2-4 binding to the decreased number of telomeric Rap1p (Alexander & Zakian, 2003) and/or the insulator activity of Tbf1p (Fourel et al., 1999). Consistent with an early loss of transcriptional silencing, we found Ade2p to be upregulated in yeast cells carrying the humanized *TLC1* at 2 scb (**Fig. 1g, TableS S11, S12**). Overall, we found the controlled loss of pAZ1 to initiate the telomeric replacement of yeast TG_1-3_ with human T_2_AG_3_ repeats and that this initiation coincided with modest proteomic remodelling. Thus we conclude that it can be used to study the long-term consequences of a telomeric repeat reprogramming.

### Chromosome end structures of humanized yeasts

We induced the telomere reprogramming by evolving 16 clones from the wild-type and humanized ancestors for 100 scb through a mutation accumulation lines (MAL) protocol, which minimizes selective pressure (**Fig. 2a, Supplementary Fig. 1 and Table S1**). During MAL, humanized yeasts’ TG_1-3_ repeats are gradually replaced by T_2_AG_3_ repeats, but they maintain a short centromere-proximal stretch of TG_1-3_ repeats (Auriche et al., 2008). Five wild-type lines became “*petites*” (growth deficiency caused by mutation or loss of mitochondrial DNA (Vowinckel et al., 2021)) and were not further characterized. We estimated that the mutation accumulation lines underwent ∼2140 and ∼1640 generations for wild-type and humanized lines, respectively (**Table S1**). Several wild-type and humanized lines subsequently were used to initiate adaptive evolution lines through 31 serial transfers (st, ∼400 generations) of large population sizes to explore adaptation to new telomeric sequences (**Fig. 2a and Supplementary Fig. 1**).

**Fig. 2.**
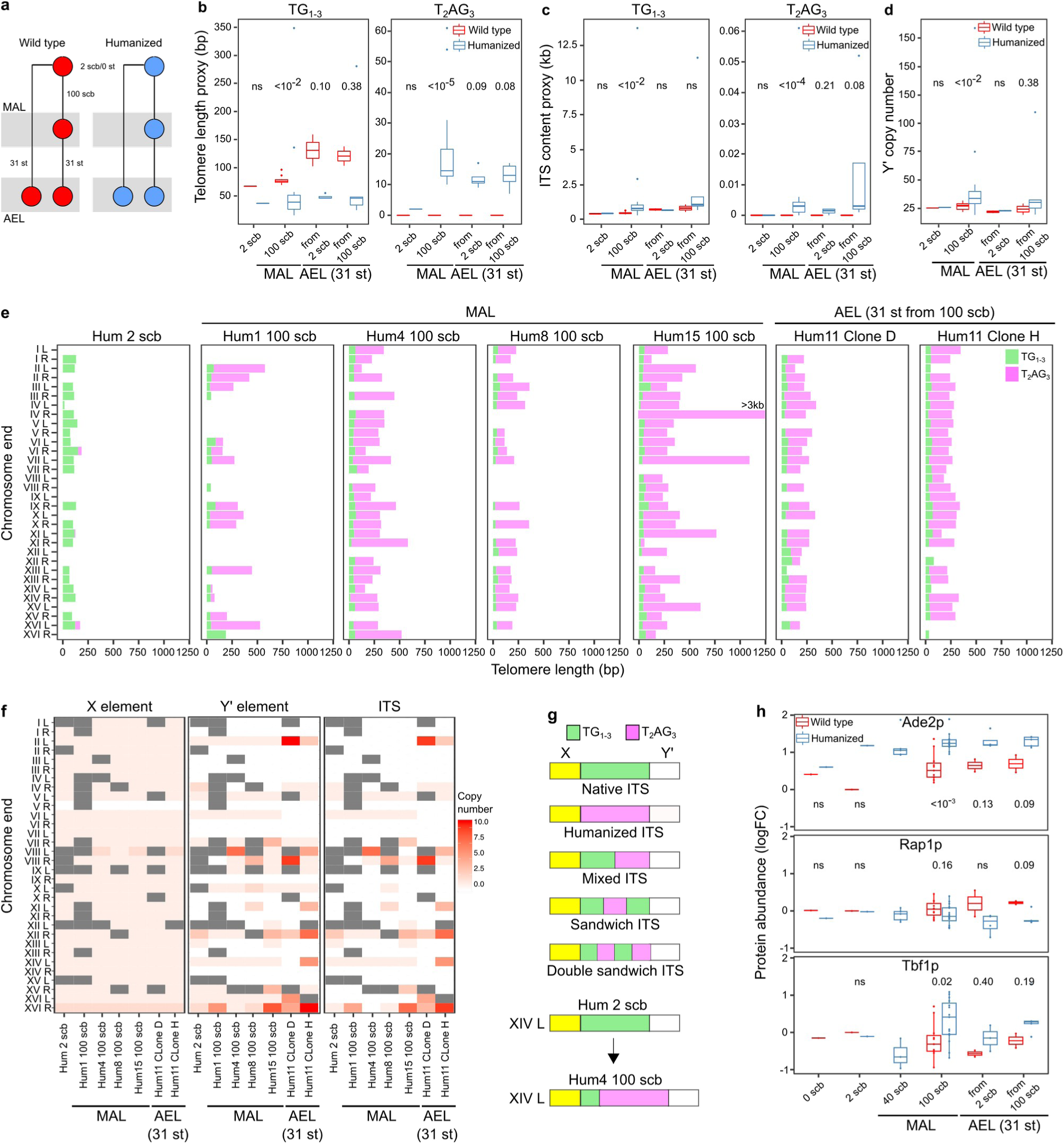
Chromosome-end dynamics during MAL and AEL. **a,** Experimental protocols used to evolve wild-type (red) and humanized (blue) ancestor strains in the absence (MAL-top panel) or in the presence (AEL-bottom panel) of selection. AEL was performed in two ways: starting from MAL at 100 scb or from the ancestors at 2 scb. **b-d** Telomere DNA length, ITS content and Y’ CN estimations in MAL at 2 (*n*=1) and 100 scb (*n*=11 and *n*=16 for wt and hum), and after AEL started from 2 (*n*=2 and *n*=4, respectively) and 100 scb (*n*=2 and *n*=5, respectively). Telomere length estimations derive from the number of telomeric reads present in short-read sequencing data. As the calculation parameters differ between TG_1-3_ and T_2_AG_3_ motifs, the respective values are not comparable between each other. Boxes: horizontal line: median, upper/lower hinge: interquartile range (IQR), whiskers: largest/smallest value within upper/lower hinge +/- 1.5x IQR. Numbers in the plots represent *p*-values resulting from comparison of wild-type and humanized lines at the underlying timepoints (two-tailed Wilcoxon test). **e**, Telomere lengths at single chromosome-ends derived from long-read genome assemblies. The order of TG_1-3_ (green) and T_2_AG_3_ (pink) repeats reflects telomere structure. Empty lines denote truncated chromosome-ends with unavailable data. In TG_1-3_-only telomeres the mapping of long reads onto the chromosome-ends showed a mixed composition comprising reads with only TG_1-3_ repeats and reads with TG_1-3_ followed by T_2_AG_3_ repeats. This underlies high heterogeneity within a line. **f**, Number of X, Y’ elements and ITS at single chromosome-ends. Grey boxes denote missing data due to truncated chromosome-ends. **g**, Upper panel: schematics of ITS structures with i) TG_1-3_-only, ii) T_2_AG_3_-only, iii) mixed ITS with a distal TG_1-3_ stretch followed by T_2_AG_3_ repeats, iv) sandwich ITS with T_2_AG_3_ repeats embedded in two TG_1-3_ stretches, v) double sandwich ITS with a repeated sequence of two mixed ITS. Double sandwich ITS are observed only in AEL. Lower panel: example of ITS that was native at 2 scb but became mixed at 100 scb. **h**, Protein abundance (log_2_FC) of Ade2p, Rap1p and Tbf1p during MAL at 0 (*n*=1), 2 (*n*=1), 40 (*n*=5 for humanized only) and 100 scb (*n*=11 and *n*=16 for wild type and humanized, respectively), and after AEL started from 2 (*n*=2 and *n*=4, respectively) and 100 scb (*n*=2 and *n*=5, respectively). Protein abundance measurements were performed in triplicate for all strains except the wild-type MAL at 100 scb (*n*=1).

To characterize the telomere DNA upon reprogramming, we applied short-read and long-read technology to both MAL and AEL. Short-read sequencing confirmed the absence of T_2_AG_3_ repeats in wild-types, while their amount increased in the humanized lines over mitotic generations (**Fig. 2b and Table S2**). ITS content and Y’ copy number (CN) increased in humanized yeasts, a hallmark of type I survivors when cells overcome telomerase inactivation via recombination-mediated telomere lengthening. Humanized yeasts also carried ITS with T_2_AG_3_ repeats suggesting telomere-subtelomere recombinations (**Fig. 2c-d and Table S2**).

Long-read sequencing can resolve telomeres at single chromosome-end resolution (Karimian et al., 2024; Sanchez et al., 2024; Schmidt et al., 2024; Sholes et al., 2022; Tham et al., 2023). We thus applied Nanopore sequencing to the humanized ancestor at 2 scb, 4 MAL and 2 single clones from the same AEL (**Fig. 2e**). At 2 scb, three telomeres (chrVI_R, chrXI_L, chrXVI_L) contained small amounts of terminal T_2_AG_3_ repeats, confirming that telomeric reprogramming starts at the first single-cell bottlenecks. At 100 scb, almost all telomeres are constituted by an internal TG_1-3_ (median 45.5±23.3 bp) and a terminal T_2_AG_3_ stretch (median 224±357.5 bp). While the TG_1-3_ length is similar across telomeres, the T_2_AG_3_ lengths show remarkable variation with the Hum15 line with ultra-long telomeres (>3 kb in chrIV_R) (**Fig. 2e, Supplementary Fig. 2a and Table S3**).

ITS and Y’ elements consistently increased in MAL and AEL, with chrVIII_L and chrXVI_R being preferential sites for their amplification (**Fig. 2f**). We confirmed the presence of T_2_AG_3_ repeats at ITS, with 5 conformations, implying recombination between telomeres and ITS (**Fig. 2g, Supplementary Fig. 2b-c and Tables S3-4**). In contrast, X elements were largely stable with 1 copy per chromosome-end, except for chrXVI_R that carried 2 copies and occasionally amplified on chrVIII_L (**Fig. 2f and Table S4**). Abundant chromosome size variations that are compatible with subtelomeric amplifications were also observed (**Supplementary Discussion 1 and Supplementary Fig. 2d**). These results raise the possibility that the amplification of subtelomeric ITS and Y’ elements compensates for the inadequate chromosome end protection in humanized yeasts. To further probe this hypothesis, we analyzed co-variation between telomere length, ITS and Y’ CN. Y’ and ITS CN were strongly correlated both in MAL and in AEL (*r*=0.96 and *r*=0.99, *p*<2.2e^-16^) (**Supplementary Fig. 2e**).

We investigated the impact of telomere humanization on proteins that directly bind telomeric repeats and the telomere silencing reporter through mass spectrometry. The abundance of Ade2p was significantly higher in humanized lines than in wild-types in both MAL and AEL, consistent with loss of transcriptional silencing in subtelomeres (*p*=0.0003 in MAL at 100 scb). This shift was mirrored by lower Rap1p levels (*p*=0.09 in AEL from 100 scb) and higher Tbf1p levels (*p*=0.02 in MAL at 100 scb), consistent with humanized yeasts adjusting the level of their telomere-binding-proteins according to telomere DNA sequence composition or length (**Fig. 2h**). In contrast, only 15 of the 147 known telomere-length-maintenance proteins that were present in our proteomics dataset showed significant difference between humanized and wild-type yeasts (**Supplementary Discussion 2**).

Overall, we disentangled the telomere and subtelomere structure of yeasts with humanized telomeres at unprecedented resolution, revealing high telomere/subtelomere dynamics and heterogeneity.

### The fitness and molecular landscape of humanized yeasts

We investigated the impact of telomere humanization on yeast fitness by measuring mitotic cell growth and chronological lifespan. During MAL, the cell doubling time remained largely stable for the wild-types, while humanized lines showed early growth defects followed by partial recovery (median doubling time 2.0/2.8 h for WT/Hum at 100 scb, *p*=1.5e^-7^). This indicates that some adaptation occurred despite the MAL being bottle-necked to random single cells (or at least extremely small populations) (**Fig. 3a, Supplementary Fig. 3a-b and Table S5**).

**Fig. 3.**
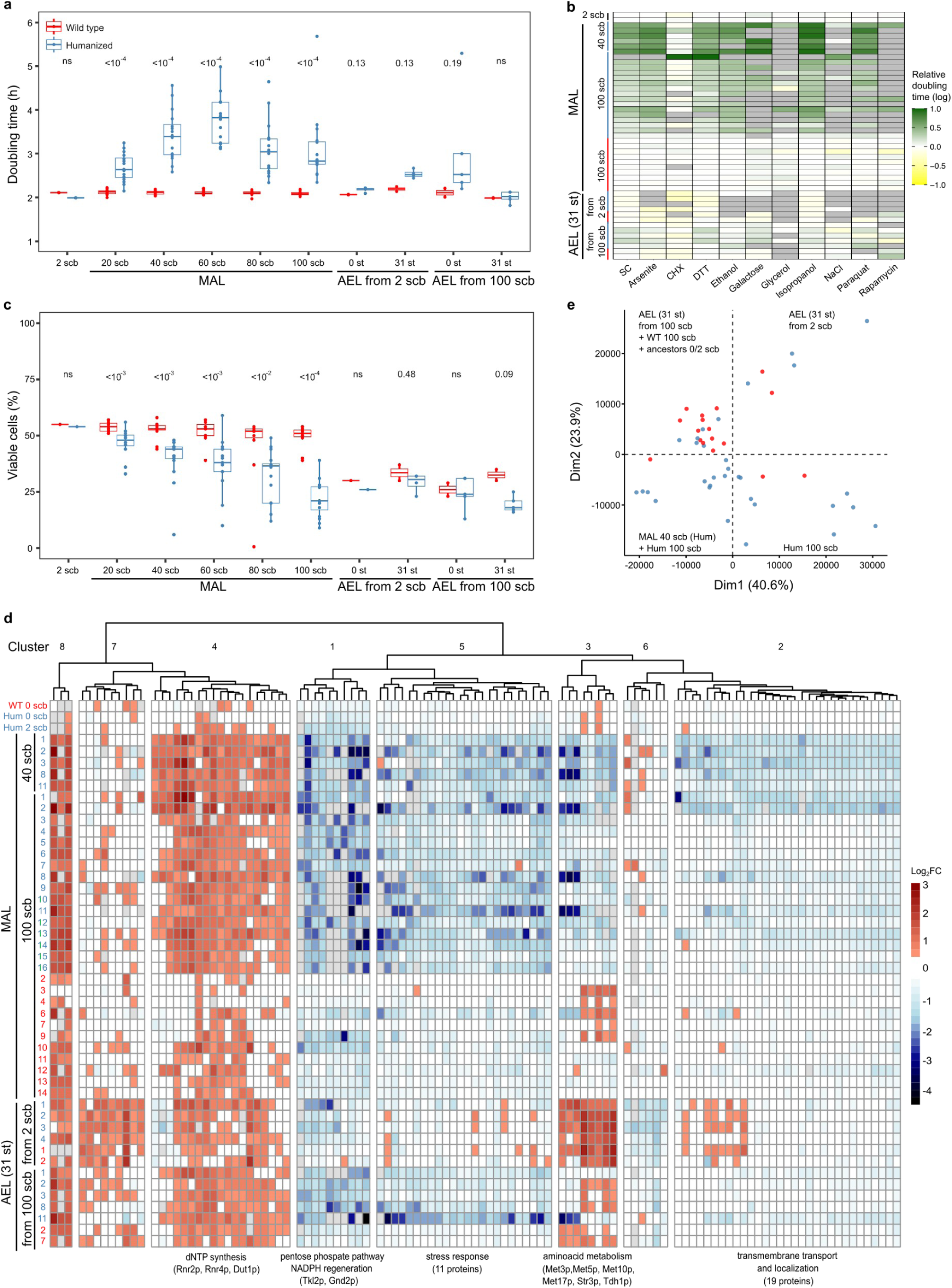
Fitness decay and recovery. **a-c,** Population doubling time, survival rate and relative doubling time of MAL every 20 scb, and of AEL. Boxplots as in Fig. 2. In Fig. **b**: green, white and yellow tones denote poor, equal or better growth respect to the control strain used for normalization (WT1 at 2 scb). Grey boxes denote missing data due to extremely poor growth. Red and blue lines denote WT and Hum yeasts, respectively. p<10^-16^ (WT vs Hum 100 scb), p <10^-6^ (WT vs Hum 100 scb/0 st), p=0.05 (WT vs Hum 100 scb/31 st), p=ns (WT vs Hum 2 scb/0 or 31 st). Fig. **c** shows values at 12 days post-saturation.Population doubling time measurements were performed in triplicate. Survival rate measurements were performed in one replicate. Relative doubling time measurements were performed in 24 replicates for SC and 12 replicates for all the other conditions. Numbers or asterisks in the plots represent *p*-values resulting from the comparison of wild-type and humanized lines at the underlying timepoints (two-tailed Wilcoxon test). \**p*<0.05, \*\**p*<0.01, \*\*\**p*<0.001, \*\*\*\**p*<0.0001. **d**, Hierarchical clustering of 110 proteins whose abundance changed (log_2_FC >1 or >-1, *p*<0.05) relative to a reference wild-type strain at 2 scb. Rows represent MA or AEL, while columns represent proteins. Upregulated and downregulated proteins are shown in red/blue, respectively (starting from log2FC >0.25 or <-0.25). Gray squares indicate missing data. Significant GO terms enrichments for the clusters and their underlying genes are indicated. Protein abundance measurements were performed in triplicate for all strains except the wild-type MAL at 100 scb (*n*=1). **e**, Principal Component Analysis based on proteomic signatures across humanized/wild-type strains at different stages of experimental evolution. Strains belonging to the 4 quadrants and separated across the two dimensions are indicated as text annotations.

We next tested whether telomere humanization impacts cell growth across environments, including different carbon sources and stresses, and observed that rapamycin and glycerol magnified the fitness defects of humanized yeasts (**Fig. 3b and Table S6**). We further measured the survival rate across a 12-days incubation in a rich medium without refreshing. Humanized lines progressively decreased their survival rate without recovering it (median viability 51/21 % for WT/Hum at day 12 and 100 scb, *p*=1.7e^-5^) (**Fig. 3c, Supplementary Fig. 3c and Table S7**).

Next, we characterized the mitotic growth of AEL started from MAL after 100 scb. During this adaptive evolution, we observed complete growth recovery (median doubling time 2.0/2.0 h for WT/Hum at 31 st, *p=*0.85). This recovery was evident also in non-adaptation environments, including rapamycin and glycerol (*p*=0.05 combining all conditions). However, it is accompanied by a decrease in survival rate (median viability 32/18 % for WT/Hum at day 12 and 31 st, *p*=0.09) (**Fig.s 3a-c, Supplementary Fig. 4a and Tables S5-7**). When started from 2 scb, both wild-type and humanized AEL had the same growth and survival and even improved their growth in certain environments when compared to the starting clones. Thus, if telomere reprogramming and selection progress in concert, cells keep their fitness (**Fig. 3a-c, Supplementary Fig. 4b and Tables S5-7**).

We examined the proteomic profiles of humanized and wild-type yeasts and derived a list of 110 proteins whose abundance changed significantly in at least one evolution condition (MAL or AEL). We used hierarchical clustering to group proteins based on their log_2_FC co-variation, uncovering 8 major clusters (**Fig. 3e**). Clusters 1,2,3,5 contain proteins down-regulated in humanized yeasts, with an enrichment for the pentose phosphate pathway/NADPH regeneration, transport/localization, aminoacid metabolism, and stress response, respectively. Proteins in clusters 1,5 remained downregulated after AEL from 100 scb, while those in clusters 2,3 returned to normal abundance. Of note, proteins in cluster 3 exhibit opposite trends between MAL and AEL from 2 scb. Cluster 4 contains proteins up-regulated in humanized yeasts and is enriched for deoxyribonucleotides synthesis. Clusters 6 and 7 comprise proteins down- or upregulated in AEL from 2 scb, respectively. Cluster 8 contains proteins upregulated in all strains, with no functional enrichment (**Fig. 3d and Table S12**). Proteomic profiles of humanized and wild-type yeasts were similar in AEL initiated from 2 scb., indicating that their differential protein abundance is mainly driven by growth advantage and faster metabolism, rather than telomere structure. This effect might underlie a common adaptation signature to the dropout medium through a modification of aminoacid metabolism. Furthermore, we performed Principal Component Analysis to group strains based on their proteomic profile and separated humanized and wild-type MAL. Humanized AEL from 100 scb clustered together with the wild type MAL and the ancestors, confirming that they recovered a proteomic profile similar to the wild types as they recovered their growth capacity. In contrast, both humanized and wild-type AEL from 2 scb clustered together (**Fig. 3e**). We further defined a “Telomere Humanization Proteomic Response” (THPR) as the proteins that are differentially abundant in at least 2 humanized MAL at 40 or 100 scb (*n*=283). We compared the THPR with genes whose RNA is differentially expressed upon telomere dysfunction induced by a *cdc13* temperature-sensitive mutation and that are present in our dataset (*n*=232), and found a significant overlap between them (*n*=79, two-tailed *X^2^* test, *p*=6.5e^-20^, **Table S12**) (Greenall et al., 2008). In contrast, we did not find a significant overlap between the THPR and Rap1p targets (*n*=345) (Brindle et al., 1990; Duveau et al., 2021; Hu et al., 2007; Lickwar et al., 2012; MacIsaac et al., 2006; McNeil et al., 1990; Song et al., 2020, Stanway et al., 1989), indicating that the THPR is not driven by Rap1p-dependent transcription (**Supplementary Discussion 2**).

Taken together, our experiments revealed that reconfiguring telomeric DNA sequences causes a severe fitness decline accompanied by a proteomic deregulation and partial reprogramming, but yeasts can recover wild-type growth.

### Mutations in the DNA damage response rescue humanized yeasts fitness

We analyzed whole-genome sequencing data from MAL and AEL to characterize the mutational spectrum and genetic alterations underlying fitness recovery. We identified 250 new single-nucleotide variants (SNVs) in the 16 humanized MAL (49/51 % within subtelomeres/core), and 87 in the 11 wild-type MAL (17/83 % within subtelomeres/core). Given that subtelomeres cover 11% of the genome (Yue et al., 2017) and assuming that selection had a negligible influence on variant distributions across MAL, we thus find that subtelomeres constitute a mutational hotspot in humanized yeasts (binomial test, *p*<0.0001). Humanized yeasts have significantly higher mutation rates than wild-types, which instead are in line with previous estimates (Sharp et al., 2018; Tattini et al., 2019) (**Fig. 4a, Supplementary Fig. 5a-c and Table S8**). A GO-term enrichment analysis revealed that wild-type MAL mutations were enriched in genes encoding ion binding proteins (*p*=0.03), while mutations in humanized MAL were enriched in genes encoding ceramid biosynthesis (*p*=0.10), cell periphery (*p*=0.06) and DNA damage response proteins (MAL Hum8 - *TEL1* and MAL Hum4 - *MEC1*; *p*=0.09), suggesting that these pathways could underlie their fitness recovery (**Fig. 4b and Table S9**). Both the *MEC1* (R1824) and *TEL1* (L2308) mutations are aminoacid substitutions located in the FAT domain of the respective proteins, with the *TEL1* mutation located in a highly conserved region and therefore predicted to be impactful. The humanized MAL carrying the *TEL1* L2308R mutation underwent massive ITS/Y’ amplifications, chromosomal rearrangements and whole-genome duplication. The *MEC1* R1824H mutation occurred between 40-60 scb whereas the *TEL1* L2308R mutation occurred between 60-80 scb. The occurrence of *TEL1* L2308R precedes fitness recovery, supporting that DDR inactivation rescues fitness in humanized yeasts (**Fig. 4c, Supplementary Fig. 3a-b and Table S5**).

**Fig. 4.**
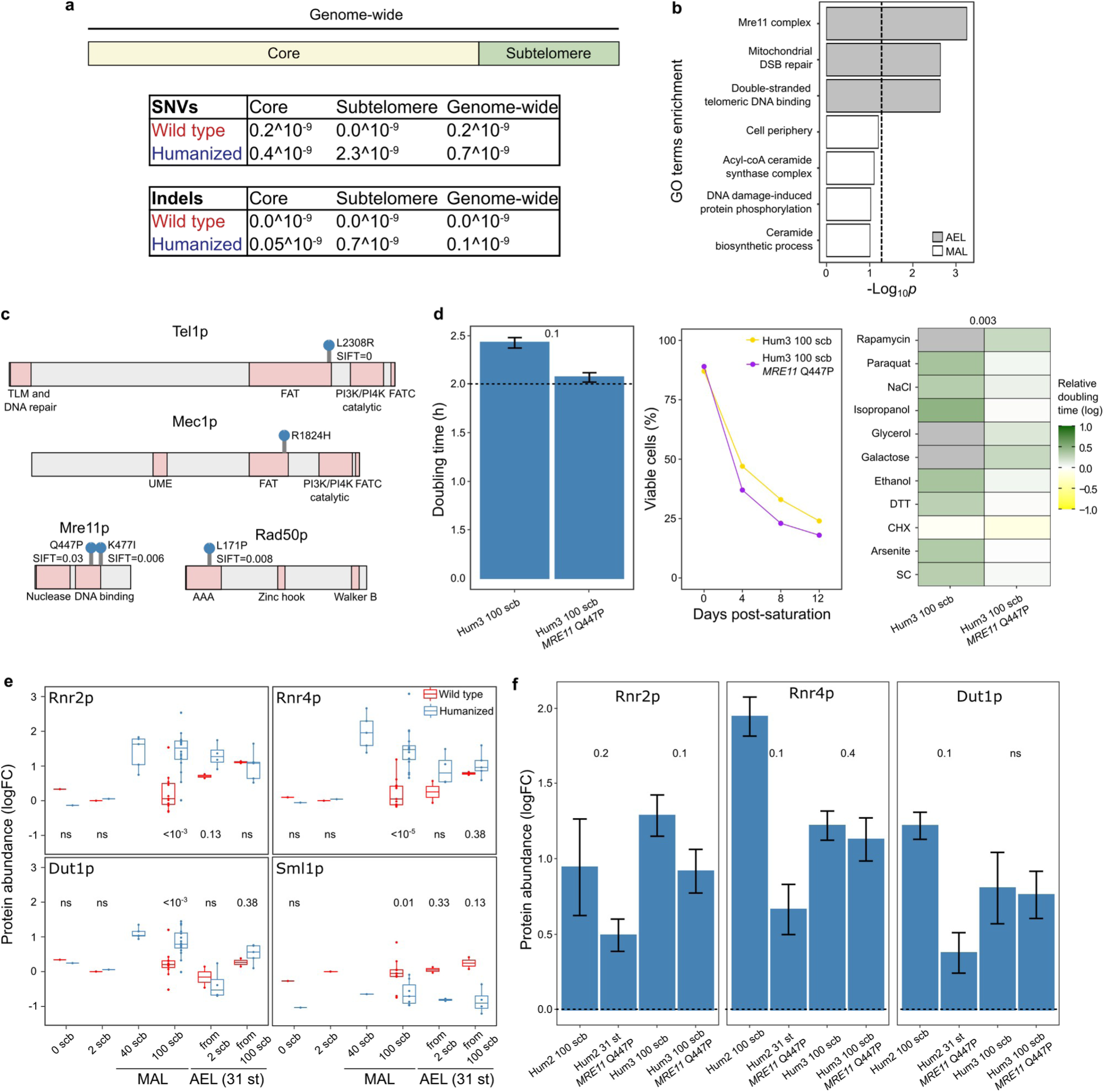
Effect of telomere humanization on mutation rate and DDR targets. **a**, Base substitution and insertion/deletion rate per bp per generation in wild-type and humanized MAL, in core (yellow), subtelomeres (green) and genome-wide. **b**, GO terms enriched for mutations occurred in humanized MAL (white bars) and AEL (grey bars). The dashed line indicates the significance threshold at *p*<0.05. **c**, Location of mutations in the DDR proteins. Protein domains are indicated by pink boxes. **d**, Population doubling time, survival rate and relative doubling time of one humanized line at 100 scb (Hum3) before and after the introduction of a missense mutation in *MRE11* (Q447P). The dashed line indicates the population doubling time of wild-type. Colour codes and number of replicates are as described in Fig. 3. *p*=0.003 for doubling time across combined conditions. Barplot: mean + standard deviation. **e-f**, Protein abundance (log_2_FC *vs* the reference wild-type at 2 scb) of Rnr2p, Rnr4p, Dut1p (**e**,**f**) and Sml1p (only in **e**) in wild-type and humanized yeasts during MAL at 0, 2 (*n*=1 for both), 40 (*n*=5 for hum only) and 100 scb (*n*=11 and *n*=16, respectively), and after AEL started from 2 (*n*=2 and *n*=4, respectively) and 100 scb (*n*=2 and *n*=5, respectively). In **f,** Sml1p has been removed due to unavailable data, and the dashed line indicates the log_2_FC of the reference. Protein abundance measurements were performed in triplicate for all strains except the wild-type MAL at 100 scb. Boxplots as in Fig. 2. Barplots: mean + standard deviation. Numbers in the plots represent *p*-values resulting from the comparison of wild-type and humanized lines from the underlying experimental evolution timepoints or genetic backgrounds (two-tailed Wilcoxon test).

Bulk sequencing of AEL at 31 st unveiled only few new mutations (*n*=13/38 in AEL starting from 2/100 scb), with the tryptophan tRNA genes recurrently affected (*n*=6). Mutations affecting components of the MRX complex, which binds DNA breaks or uncapped telomeres and then activates the DDR through the recruitment of Tel1p and Mec1p, occurred recurrently and independently in three AEL: Hum1 - *MRE11 K477I*; Hum2 - *MRE11 Q447P*; Hum3 - *RAD50 L171P*). These lines grew poorly at 0 st but completely recovered their fitness after AEL (31 st), and all three mutations are predicted impactful through SIFT score analysis (**Fig. 4c**). The two *MRE11* mutations in the DNA binding domain and *RAD50* mutation in the ATPase domain were all fixed, in line with that they are strongly beneficial (**Fig. 4b-c and Tables S8-9**). To confirm the adaptive role of the MRX mutations, we engineered *MRE11* Q447P into the slow-growing humanized line 3 at 100 scb. This mutation produced a full growth recovery across conditions, but decreased the survival rate (**Fig. 4d and Tables S5-7**). We tested whether the beneficial effect would also manifest in fit humanized yeast that had not yet suffered a loss of reproductive capacity by introducing *MRE11* Q447P in the humanized line 1 at 2 scb. However, we observed no improvement in growth or lifespan, consistent with that its benefits are strictly limited to compensating for the fitness loss arising from telomere reprogramming (**Supplementary Fig. 5d and Tables S5-7**). The full deletion of *MRE11* improved both growth and survival rate but did not match the growth of the wild type MAL (**Supplementary Fig. 5e and Tables S5-7**). Taken together, these results suggest that *MRE11* mutations induce partial loss-of-function.

To further disentangle how DDR modulation rescues fitness in humanized yeasts, we inspected protein abundance of known DDR targets in the proteomic dataset. We observed upregulation of the ribonucleotide reductase (RNR) subunits Rnr2p/Rnr4p and the deoxyuridine triphosphate diphosphatase Dut1p, and the downregulation of the RNR inhibitor Sml1p. The upregulation of Rnr2p/Rnr4p/Dut1p is highest in MAL at 40 scb while it decreases after AEL from 100 scb (*p*<10^-2^ for all proteins at 100 scb) (**Fig. 4e**). We further checked the abundance of Rnr2p/Rnr4p/Dut1p in a humanized AEL that spontaneously developed a mutation in *MRE11* (Hum2 31 st *MRE11* Q447P), as well as a line where we introduced the same mutation by CRISPR/Cas9 genetic engineering (Hum3 100 scb *MRE11* Q447P). The three proteins decrease when *MRE11* Q447P is present, suggesting attenuation of the DDR (**Fig. 4f**).

Overall, these findings reveal that humanization of yeast telomeres increases mutation rate and the partial inactivation of DDR rescues their fitness through its downstream targets.

### Chromosome XVI aneuploidy is adaptive in humanized yeasts

After revealing the beneficial effect of DDR point mutations, we investigated whether adaptation also occurred via large-scale copy number and structural variants (SVs). All the MAL remained haploid except for a whole-genome duplication in the humanized MAL 8 with a *TEL1* loss-of-function mutation. This line likely underwent endoreduplication since the *MAT* locus remained homozygous (*MATa/MATa*) (**Fig. 5a and Table S10**) (Harari et al., 2018). We next analyzed long-read-sequencing-derived genome assemblies from 4 humanized MAL at 100 scb (Hum1,4,8,15) and an ancestor at 2 scb. We did not detect any major SVs except for Hum8 at 100 scb, which underwent a translocation between chromosomes V and XV mediated by the two *ADE2* copies (**Supplementary Fig. 6, Table S10 and Supplementary Discussion 1)**.

**Fig. 5.**
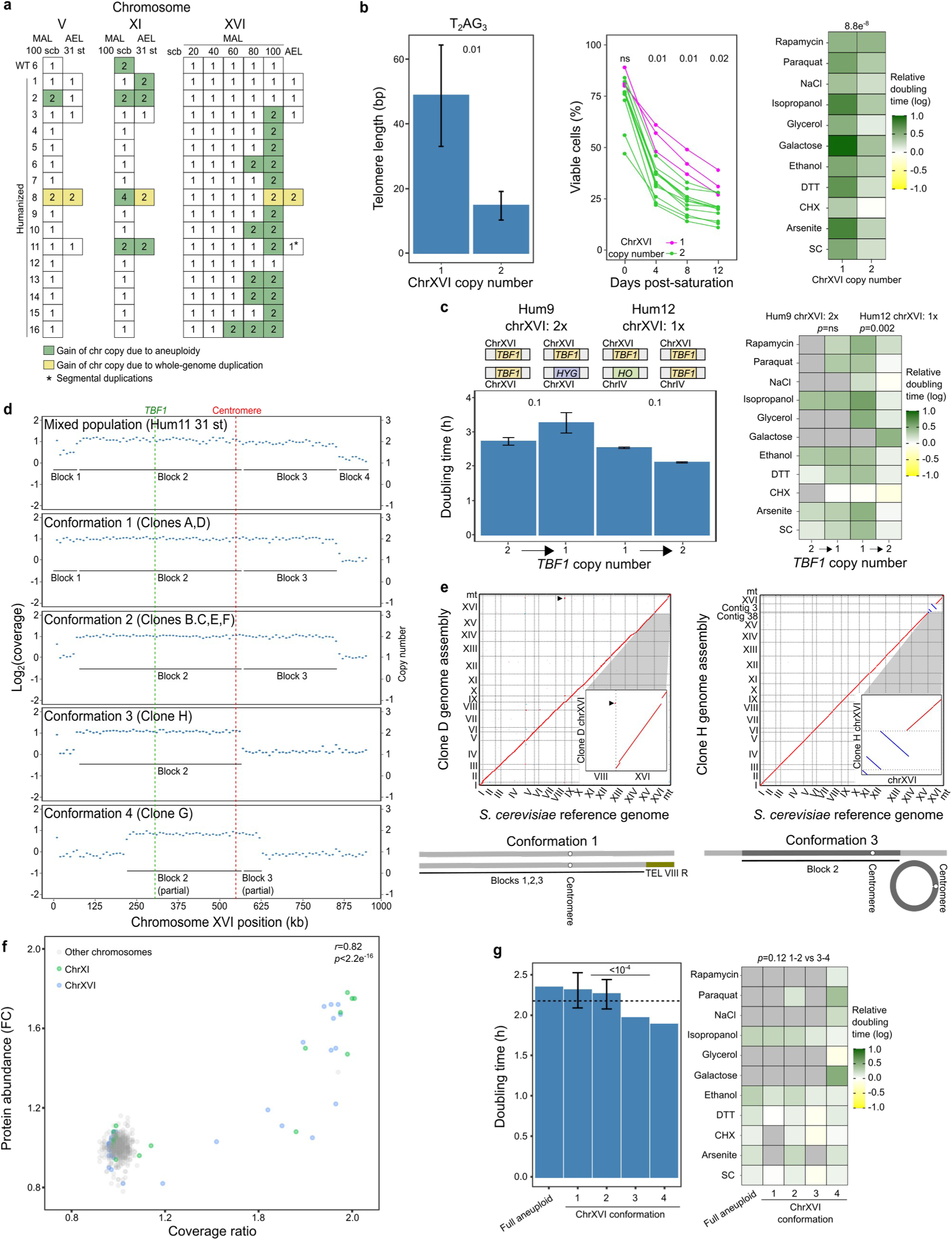
Structural variation during MAL and AEL. **a**, Chromosome-scale copy number of chromosomes that underwent copy number variation in the MAL and AEL. White boxes represent euploid chromosomes that maintained their original copy number (1), while green/yellow boxes indicate gain of copy by aneuploidy or whole-genome duplication, respectively. **b**, T_2_AG_3_ telomere length, survival rate and relative doubling time in humanized MAL that are either euploid (one copy) or aneuploid (two copies) for chromosome XVI. Barplot: mean + standard deviation. Color codes and number of replicates are as described in Fig. 3. Hum8 was excluded due to its diploidy. Numbers in the plots represent *p*-values resulting from the comparison of humanized lines from the underlying genetic backgrounds (two-tailed Wilcoxon test). **c**, Population and relative doubling time of humanized lines that underwent the knock-out (left) or the knock-in of one copy of *TBF1*, before and after the genetic engineering. The arrow indicates the direction of the genetic engineering. Numbers in the plots represent *p*-values resulting from the comparison of humanized lines from the underlying genetic backgrounds (two-tailed Wilcoxon test). Barplot: mean + standard deviation. **d**, Coverage profiles on chromosome XVI for an AEL started from 100 scb (top panel) and of 8 single clones derived from it (bottom panels). Dashed lines indicate the position of *TBF1* (green) and the centromere (red). Plain lines indicate the length of duplicated blocks as defined by the points of coverage breakage in the mixed population (top panel). **e**, Comparison of the *S. cerevisiae* reference genome (x axis) and our Nanopore-derived genome assemblies for two clones (D,H) derived from the mixed population. Sequence homology signals are indicated in red (forward match) or blue (reverse match). Insets show a zoom-in of chromosome XVI. Karyotypes on the right show the structure of the two chromosome XVI copies. **f**, Correlation between chromosome copy number estimated from short-read sequencing coverage (x axis) or from normalized FC (y axis), across samples for whom we possessed both sequencing and proteomics data (*n*=44). Aneuploid chromosomes (XI and XVI) are highlighted in colour. **g**, Population and relative doubling time of clones grouped by their chrXVI conformation (*n*=2,4,1,1 for conformation 1,2,3,4 respectively). The fully aneuploid (for chrXVI) Hum11 at 100 scb is shown on the left (*n*=1). The dashed line indicates the doubling time of the mixed population (Hum11 at 31 st). Numbers in the plots represent *p*-values resulting from the comparison of humanized lines from the underlying conformations (two-tailed Wilcoxon test). Barplot: mean + standard deviation.

We next analysed read coverage along chromosomes to determine whether large CNV and aneuploidies occurred during MAL. We discovered one segmental duplication of chromosome IV (∼100 kb, WT2) and 17 whole chromosome aneuploidies in our lines. These were all chromosome gains and exclusively occured in humanized lines (except one in WT6). Chromosome XVI was aneuploid in 12 lines suggesting a possible adaptive role (**Fig. 5a and Table S10**). We further explored whether an additional chromosome XVI is beneficial by comparing humanized lines carrying 1 or 2 copies of this chromosome. Lines with 2 copies had shorter telomeres, and this difference was driven by fewer T_2_AG_3_ repeats (*p*=0.01) (**Fig. 5b, Supplementary Fig. 7a and Table S2**). Furthermore, they had a shorter doubling time in multiple environments but lower survival rate (*p*=8.8e^-8^ for doubling time across conditions, *p*=0.02 for survival rate at day 12) (**Fig. 5b, Supplementary Fig. 7b and Tables S5-7)**. To better pinpoint the emergence of the aneuploidy during MAL evolution, we probed its existence in the stored records through qPCR and found that it occurred between 60-100 scb (only one case at 40-60 scb), with its emergence correlating well with the decrease of both doubling time and survival rate, confirming a causative role (*p*=0.22/0.02 for doubling time/survival at day 12) (**Fig. 5a, Supplementary Fig. 7c-e and Tables S5-7**).

Aneuploidies are usually detrimental in nature, especially when involving long chromosomes, but can also fuel adaptation (Pompei & Cosentino Lagomarsino, 2023; Rojas et al., 2024; Sheltzer & Amon, 2011). We inspected whether any telomere-binding-proteins were encoded on chromosome XVI, detecting Tbf1p, which directly binds T _2_AG_3_ repeats (Brigati et al., 1993). We tested the effect of *TBF1* duplication by engineering its copy number, first by deleting one *TBF1* copy in a humanized MAL carrying two chromosomes XVI (Hum9) and then by inserting a second *TBF1* copy in a humanized MAL carrying a single chromosome XVI (Hum12). We found that the addition of a second *TBF1* copy to a humanized MAL carrying one improved growth, while the deletion of one out of two *TBF1* copies increased the doubling time by 17% (**Fig. 5c**). The beneficial effect of an extra *TBF1* in humanized yeasts likely derives from better telomere capping and/or from Tbf1p’s inhibitory action towards the binding of the MRX complex to telomeres and its subsequent activation of DDR, as supported in our proteomic data by the inverse relationship between protein levels of *TBF1* and DDR targets (Rnr2p, Rnr4p, Dut1p) (Berthiau et al., 2006; Fukunaga et al., 2012) (**Supplementary Fig. 8a-b**). These results suggest that the fitness advantage conferred by *TBF1* duplication is due to a combination of telomere protection and DDR-inhibitory functions.

Next, we investigated the evolutionary dynamics of large-scale structural variants during AEL. All the AEL derived from ancestors at 2 scb, i.e. from before substantial telomere reprogramming, maintained their original ploidy and showed no major structural variants. In contrast, AEL derived from MAL at 100 scb and that had gained extra chromosomes during the MAL evolution, frequently lost the extra chromosomes during the adaptive evolution and thereby restored a balanced genome content. Humanized lines 2,8,3 lost their additional copies of chromosomes V, XI and XVI, respectively, while Hum11 revealed a mixed population of clones carrying different segments of the extra copy of chromosome XVI (**Fig. 5a and Table S10**). We isolated 8 single clones from this population and uncovered 4 distinct chromosome XVI conformations by qPCR and whole-genome sequencing. All the segments’ breakpoints were flanked by repetitive sequences such as Ty and long terminal repeats that likely mediated the recombination events. Notably, the region encompassing the *TBF1* gene remained duplicated in all conformations and included the centromere (CEN16) (**Fig. 5d**). We applied long-read sequencing to two of these clones to disentangle their chromosome XVI structure. Clone D carried almost the full version of duplicated chromosome XVI, except for a terminal segment, and the duplicated segment was stabilized by the duplication and translocation of chrVIII-R telomeres. Clone H carried only a central duplicated segment of chromosome XVI and this segment was circularized with the circularization likely mediated by Ty elements and stabilized by CEN16 (**Fig. 5e**). This circular structure is likely present in all conformations lacking terminal duplicated blocks (i.e. conformations 2,3,4) and has the advantage of not needing telomeres. We tested whether these segmental duplications resulted in a double amount of proteins. Proteomic profiles mirrored read coverage ones for the 4 chrXVI conformations (**Supplementary Fig. 8c**). However, the average abundance for proteins encoded by genes in these segments was less than 2-fold higher than that of other proteins, indicating a buffering effect. An extension of this analysis to the whole genome revealed a partial gene dosage compensation at the proteome level, which was probably selected during the long-term evolution (**Fig. 5f and Supplementary Discussion 3**) (Muenzner et al., 2024). We reasoned that this conformations mixture might be the result of selection favouring conformations carrying the shortest duplicated block containing *TBF1*, in order to minimize the detrimental effects associated with the duplication of all chrXVI genes. To test this hypothesis we measured growth in the 8 clones and confirmed an inverse relationship between growth and chrXVI duplicated content (*p*=2.9e^-5^ between conformations 1-2 and 3-4) (**Fig. 5g and Tables S5-7**).

Taken together, these results show that chrXVI aneuploidy is a primary evolutionary route to fitness recovery driven by the duplication of *TBF1*.

## Discussion

In this work, we studied the evolution of budding yeast cells carrying humanized T _2_AG_3_ telomeric repeats through the combination of two laboratory evolution protocols: mutation accumulation lines (MAL) and adaptive evolution lines (AEL). Humanizing the telomeric repeats opens the possibility to investigate the long-term impact of telomeric reprogramming enabling adaptation via the “pre-adapted” factor Tbf1p, in contrast to other random unnatural telomeric repeats whose usage will require long-term evolution of other proteins that will be outcompeted by the emergence of telomerase-negative mechanisms. Our results show that humanized yeasts accumulate T_2_AG_3_ repeats, and that this telomere sequence reprogramming impairs fitness and increases genome instability. This is in agreement with humanized telomeres inducing replication fork stalling and cell cycle delays, and being subject to a high repeat turnover (Bah et al., 2004; Bah et al., 2011; Di Domenico et al., 2009), thus explaining the need for massive subtelomeric amplifications of ITS and Y’ elements to prevent chromosome-end uncapping. Increased genome instability as a consequence of telomere dysfunction was already observed in humanized yeasts (Di Domenico et al., 2013) and in telomerase-deficient *est1Δ S. cerevisiae* strains (Hackett et al., 2001). However, this effect was restricted to chromosome-end regions, while our results show that telomere dysfunction increases mutation rates across the whole genome.

After an initial decline, humanized yeasts recovered their fitness through two evolutionary routes: by acquiring mutations in components of the DNA damage response or by increasing the copy number of telomere-binding proteins. The mutations in DDR genes are likely partial or complete loss-of-functions, as their appearance coincides with a decreased expression of DDR targets like *RNR* genes, suggesting a release of cell-cycle blockage induced by telomere uncapping. Previous studies showed that DDR is mildly but chronically activated in humanized yeasts and causes a G_2_/M delay (Di Domenico et al., 2009), while the deletion of the DDR mediator *RAD9* abolished DDR and the cell cycle delay (Bah et al., 2011), in line with our findings. While DDR inactivation relieves the cell cycle blockage and rescues growth, it also increases genome instability. Our humanized line 8, carrying a loss-of-function mutation in *TEL1*, exemplifies this tradeoff, as the appearance of the mutation is followed by complete growth recovery coupled with multiple chromosomal rearrangements, extensive subtelomeric amplifications and even whole-genome duplication. Given its important role in DNA repair, DDR inactivation likely constitutes a temporary solution while its long-term inactivation would probably lead to genomic melt-down. A more likely scenario of telomere sequence evolution might involve the transient inactivation of the DDR followed by the occurrence of compensatory mutations in telomere-binding proteins to adapt to their novel telomeric sequences.

The second route that humanized yeasts use to rescue their fitness passes through the duplication of chromosome XVI, ultimately to increase the abundance of Tbf1p, a T _2_AG_3_ binding protein (Brigati et al., 1993). While aneuploidy is generally considered detrimental, it can be beneficial under different environments (Gilchrist & Stelkens, 2019; Pavelka et al., 2010; Rancati et al., 2008). Our results are consistent with aneuploidy as a faster strategy to recover fitness, due to the high chromosome duplication rate (Yona et al., 2012). Even though MAL are generally considered a close-to-neutral evolutionary scenario, adaptive aneuploidies have been previously observed in these settings, e.g. chromosome XII is frequently duplicated in MAL propagated under rapamycin treatment to maintain a balanced rDNA content after rapamycin-induced rDNA repeat contraction (Li et al., 2023). However, aneuploidy also entails a fitness cost due to the indiscriminate increase in expression of many genes and the stoichiometric imbalance of proteins involved in protein complexes (Papp et al., 2003). Thus, the ultimate evolutionary outcome is dictated by a tradeoff between the fitness costs and gains of the aneuploidy. Removing the parts of the duplicated chromosome that are not driving the fitness improvement constitutes a successful strategy, as exemplified in one of our AEL where the clones with the shortest duplicated chromosome XVI segments were the fittest. The formation of extrachromosomal circles carrying duplicated genes, as well as other chromosomal rearrangements, has already been observed both in laboratory evolution and telomerase negative mutants (Liti & Louis, 2003; Hull et al., 2017; Libuda & Winston, 2006). Our results provide a temporal dynamic to the recent observation of complex aneuploidies associated with structural variations, showing that aneuploidies precede structural variations (O’Donnell et al., 2023). These results are also in line with a theoretical framework in which the fitness cost of an aneuploidy is directly proportional to the number of genes located on that chromosome and it is counterbalanced by the fitness benefit resulting from the dosage increase of specific genes (e.g. *TBF1*) (Pompei & Cosentino Lagomarsino, 2023). DDR mutations and *TBF1* duplication might ultimately impinge on the same outcome: the attenuation of the DDR and release of cell cycle blockage. In fact, the MRX complex binds critically short telomeres, recruits telomerase and activates the DDR through the downstream action of Tel1p. Artificially placing Tbf1p-bound T_2_AG_3_ repeats in subtelomeric regions shortens the respective telomeres, as Tbf1p inhibits MRX binding of telomeres, thus preventing the initiation of the DDR, cell cycle blockage and telomerase recruitment (Berthiau et al., 2006; Fukunaga et al., 2012; Ribaud et al., 2012). The inhibition of telomerase recruitment by Tbf1p also explains the shorter T_2_AG_3_ tracts detected in humanized yeasts carrying two copies of chromosome XVI.

Overall, our results point to an evolutionary scenario in which aneuploidy of chromosome XVI and amplification of Tbf1p is the first adaptation to the modification of telomeric sequences, due to its high frequency. However, given its fitness cost, it is removed as soon as less burdening point mutations in the DDR and MRX complex appear. In contrast, if the adaptive point mutations do not occur, another strategy is to remove the parts of the duplicated chromosome XVI that are not immediately useful, only retaining duplicated segments encompassing the *TBF1* gene. This scenario is in line with previous research showing that aneuploidy and copy number variations are quick responses to environmental stress whose role is to give time to the cells to develop more specific and losg-lasting genetic changes in the form of point mutations (Yona et al., 2015).

T_2_AG_3_ repeats are thought to be the ancestral ones from which all the others evolved (Fulnečková et al., 2013). Going back to an ancestral state might be easier than adapting to completely new challenges. This view is consistent with the fact that Tbf1p is functionally related to the telomere protective proteins TRFs in animals, and particularly in its DNA binding domain interacting specifically with T_2_AG_3_ repeats (Bilaud et al., 1996). Therefore, yeasts might behave differently if evolved with other telomeric sequences and in different environmental conditions. Mutations in the telomerase RNA template cause growth defects and telomere length dysregulation (Lin et al., 2004; McEachern & Blackburn, 1995), but it is not known how these strains would survive and adapt when propagated under selection for a long time. Recently, a variant telomeric repeat has been identified in a human family and shown to be inherited across one generation, indicating that telomere sequence modification is possible even in humans (Hinchie et al., 2024). Moreover, telomeric sequences can be introgressed from other species following horizontal gene transfer events, as exemplified by the discovery of mixed telomeric sequences in *S. cerevisiae* natural isolates, constituted by centromere-proximal *Torulaspora* telomeric repeats followed by native TG_1-3_ telomeric repeats (O’ Donnell et al., 2023). Future studies investigating the long-term impact of other telomeric repeats, as well as their interactions with the external environment, will further elucidate how cells can evolve dysfunction in key cellular processes.

## Acknowledgements

We thank Simone Mozzachiodi for help with CRISPR/Cas9 genetic engineering, Jing Li and Jia-Xing Yue for help in generating and analyzing long-read sequencing data, and Daniela Ludwig, Kathrin Textoris-Taube and Michael Mülleder at the Core Facility - High Throughput Mass Spectrometry, Charité University Medicine, Berlin, for help with sample preparation and mass spectrometry analysis. We also thank Miguel Ferreira and Zhou Xu for critical reading of the manuscript. This study was supported by ANR (ANR-18-CE12-0004, ANR-20-CE12-0020, ANR-23-CE12-0031, ANR-24-CE12-7740), Fondation pour la Recherche Médicale (EQU202003010413), Fondation ARC (No. PJA32020070002320), Impulscience - Fondation Bettencourt Schueller to G.L, VR-2022-03024 to JW and the European Research Council (ERC) under grant agreement ERC-SyG-2020 951475 to M.R. M.D. was supported by PhD fellowships from the LABEX SIGNALIFE ANR-11-LABX-0028-01 and the Fondation pour la Recherche Médicale (FDT201904008453).

## Contributions

M.D., B.B., E.G. and G.L. designed the experiments; M.D. performed the experiments and the genomic analyses; M.D.C. estimated copy number variants from short-read sequencing data; N.T. analyzed *TLC1* variation data; J.H. and M.D. generated and analysed the growth phenotype data; J.M., S.K. and M.D. generated and analysed proteomics data; J.W., M.R., E.G. and G.L. contributed with resources and reagents; E.G. and G.L. conceived and supervised the project; G.L. coordinated the project; M.D. and G.L. wrote the paper.

## Data availability

The genome sequencing generated in this study are available at Sequence Read Archive (SRA), NCBI under accession codes BioProject ID PRJNA985049, Biosample ID SAMN35786841-SAMN35786899. The phenotyping and proteomics data are available at: https://github.com/mdangiolo89/Humanized_yeasts_project and within the supplementary information files. All the strains generated in this work are available upon request. Raw and processed mass spectrometry data have been deposited at the ProteomeXchange consortium via the PRIDE partner repository with the dataset identifier PXD064562 and will be publicly released upon publication of the study.

## Code availability

YeaISTY (Yeast ITS, Telomeres and Y′ elements estimator) and other custom scripts are available at: https://github.com/mdangiolo89/Humanized_yeasts_project.

## Materials and methods

### Yeast strains and plasmids

The two ancestor strains (one for the humanized and one for the wild-type lines) derive from (Bah et al., 2004) and their genetic background is: RWY12 (BY4741 *Mat*a, *ura3-52*, *lys2–801*, *ade2-101*, *trp1*-Δ*1*, *his3*-Δ*200*, *leu2*-Δ*1*, *tlc1*Δ*::LEU2,* VR-*ADE2*-T) (Dionne & Wellinger, 1996). These strains are haploid and auxotrophic for uracil, lysin, tryptophan and histidine. The native *ADE2* gene on chromosome XV is not functional and another copy has been inserted into the right subtelomere of chromosome V. *ADE2* inactivation results in red colonies through the accumulation of purine precursors in the vacuole and can be used as a marker of transcriptional silencing at subtelomeric sites (Zonneveld & Van Der Zanden, 1995). The replacement of the *TLC1* gene with an auxotrophic marker ensures that the strains cannot use their native RNA template to elongate telomeres.

In addition, the two ancestors contain two centromeric plasmids: pAZ1 and p *TLC1*. The pAZ1 plasmid derives from (Dionne & Wellinger, 1996) and contains the wild-type version of *TLC1* and the selection marker *URA3*. The p*TLC1* plasmid backbone originally derives from pRS314 in (Sikorski & Hieter, 1989) and contains the *TLC1* gene and the selection marker *TRP1*. The *TLC1* gene in the p*TLC1* plasmid is different in the two ancestors: the wild-type ancestor carries the native p*TLC1*-y gene while the humanized ancestor carries a modified p*TLC1*-h gene in which the 16 template nucleotides (5’-CACCACACCCACACAC-3’) have been replaced by a humanized counterpart (5’-CUAACCCU-3’).

### Mutation accumulation lines

The humanized and wild-type ancestors were revived from frozen stocks and plated on Leucin/Tryptophan/Uracil dropout medium to select for cells containing both the pAZ1 and p*TLC1* plasmids. Plates were incubated for 3 days at 30 °C. After that, we streaked for single colonies on 5-FOA plates to retrieve cells that had lost the *URA3*-containing pAZ1 plasmid, and therefore rely only on the *TLC1* copy present in the p*TLC1*-y/p*TLC1*-h plasmids. We selected 16 colonies derived from the wild-type ancestor and 16 derived from the humanized ancestor, which contained only the p*TLC1*-y and p*TLC1*-h plasmids, respectively. Since two streaking-for-single were performed during the pAZ1 exclusion procedure, we dated this timepoint as 2 single cell bottlenecks (scb).

We propagated the 32 lines on Leucin/Tryptophan dropout solid medium in order to maintain cells with *tlc1::LEU2* genotype and to retain the p*TLC1*-y/p*TLC1* at the final timepoint (100 scb) and at four intermediate timepoints (20, 40, 60, 80 scb) were stored at -80 °C. Moreover, we stored 4 humanized and 2 wild-type lines at 2 scb, immediately after the loss of pAZ1.

To estimate the number of generations occurring every 72 h, three colonies derived from three independent humanized lines at every timepoint and one colony from a wild-type line at 2 and 100 scb were resuspended in 100 µl of sterile water and serially diluted. Twenty µl of each dilution were plated on solid YPD medium and incubated for 72 h at 30 °C. The number of colonies was manually counted in the plate with a suitable dilution and the number of generations (*G*) was estimated according to *G*=log_2_(*n*×*d*), where *n* is the number of cells on the plate and *d* is the corresponding dilution factor. The total number of generations occurred during the MAL protocol for the wild-type and humanized lines (*T_wt_*and *T_hum_*) were calculated as: *T_wt_=G×100* where *G* was averaged across the only representative line at 2 and 100 scb; *T_hum_=(G_20_×20)+(G_40_×20)+(G_60_×20)+(G_80_×20)+ (G_100_×20)* where *G_x_* was averaged across the three representative lines at each timepoint. The humanized lines at 2 scb were not considered for the calculation.

### Adaptive evolution

The 4 humanized and 2 wild-type lines at 2 scb, as well as 5 humanized lines and 2 wild-type lines derived from the MAL protocol (Hum1,2,3,8,11,WT2,7) were further submitted to adaptive evolution (AEL) by multiple serial transfers (STs) of large population sizes. Strains were inoculated in tubes containing 5 ml of Leucin/Tryptophan dropout liquid medium and incubated at 30 °C in a shaking incubator. Serial transfers were performed every 48h by diluting 50 μl of the culture into 5 ml of fresh Leucin/Tryptophan dropout medium (1:100 dilution). We performed a total of 31 serial transfers.

### Estimation of the growth performance in liquid medium

Strains were incubated two nights in 200 uL of liquid Leucin/Tryptophan dropout medium in a non-shaking incubator at 30 °C. The day after, strains were mixed and 2 ul of the mixed culture was resuspended in 200 uL of fresh Leucin/Tryptophan dropout liquid medium in a 96-well plate. The plate was incubated in a TECAN plate reader machine (Tecan, Infinite F200 Pro) at 30 °C for 72 hours and optical density (600 nm) was measured every 15 minutes during the experiment. Resulting data were analyzed using the software PRECOG (Fernandez-Ricaud et al., 2016). We phenotyped 3 biological replicates for each strain. Given the high number of strains in this study, phenotyping was performed using several 96-well plates. To correct for the batch effect and compare data across plates, we included the humanized ancestor at 0 scb (MJD548) in all the plates and used it as internal control. We normalized the data in each plate by dividing by the average of the 3 control replicates in that plate. We then recalibrated all the measurements by multiplying for constant values: 1.97 for population doubling time, 5.82 for lag duration and 3.20 for yield. These values represent the average of the 3 control replicates in the first plate. Values reported in Table S5 and in the Fig.s are already recalibrated.

### Estimation of the growth performance across multiple conditions

Strains were grown in 5 ml Synthetic Complete (SC) shaking liquid culture overnight. 100 uL from each of these cultures were resuspended into four 96-well plates, each with a different randomized layout. These four 96-well plates were then transferred onto a 1536 SC PLOS agar plate using the Rotor HDA Robot (Singer ltd). Each randomized layout was replicated 3 times onto the 1536 plate, resulting in a total of 12 replicates (24 for SC). Each plate also contained 384 positions with spatial controls (Zackrisson et al., 2016). The spatial control used was a wild-type ancestor at 2 scb (MJD5). Due to the size differences of the colonies when transferring from liquid to solid, the 1536 solid plate was grown for 3 days and then replica-plated onto a new SC agar plate. This new plate is the preculture plate. After 3 days of growth on the preculture plate, the strains were transferred onto the plates with the different stressors and phenotyped. Lag duration, population doubling time and yield were extracted from the resulting growth curves using Scanomatic (version 2.2). Lag was calculated as the intercept between the tangent of the initial colony size (average of the first three timepoints) and the tangent from the timepoints used to calculate the growth rate. Population doubling time was calculated at the steepest incline of the growth curve, using five timepoints for a local regression. Yield was calculated as the difference between the initial colony size, as calculated for lag, and the final colony size (average of the last three timepoints). All measurements were calculated from the smoothened growth curves.

### Estimation of survival rate

Strains were grown two nights in liquid Leucin/Tryptophan dropout medium, diluted 100x in 200 μL of fresh Leucin/Tryptophan dropout in a 96-well plate and incubated at 30 °C for 15 days. We measured viability at the beginning of the stationary phase when cultures reached saturation (3 days), and then every 4 days until day 15. Data are shown by days post-saturation, meaning that day 0 post-saturation corresponds to day 3 after the inoculation of the culture, while day 12 post-saturation corresponds to day 15 post-inoculation. At each time point, 5 μL of cells were transferred in 100 μL of staining solution (Phosphate-buffered saline-PBS + 3 μM propidium iodide + 200 nM YO-PRO-1) in a 96-well plate and incubated 10 minutes in the dark at 30 °C. Cell viability was measured by high-throughput flow cytometry on a Cytoflex machine, using the plate reader module. Cells were excited with the 561nm Yellow/Green laser. Fluorescence is read with the Y610-mCHERRY filter for propidium iodide and with the B610-ECDA filter for YO-PRO-1. Propidium iodide and YO-PRO-1 are membrane-impermeable nucleic acid binding molecules that can only enter into dead cells whose cellular membrane is damaged. Non-fluorescent cells were counted as viable whereas fluorescent cells were counted as dead. We measured one biological replicate with 3000 events per sample. Flow cytometry data were subsequently analysed using the CytExpert software.

### Ploidy estimation and pulsed-field gel electrophoresis

Ploidy was estimated for MAL at 2 and 100 scb and after AEL (at 31 st) using a propidium iodide (PI) staining assay. Ploidy was estimated at all timepoints during the MAL protocol for humanized line 8, which showed a diploid genome content at 100 scb. Cells were incubated in 100 uL of Leucin/Tryptophan dropout liquid medium in a 96-well plate and left two nights at 30 °C. After that, 3 μl were incubated in 100 μl of cold 70% ethanol for 3 h at 4 °C, washed twice with PBS, and resuspended in 100 μl of staining solution (15 μM PI, 100 μg/ml RNase A, 0.1% v/v Triton-X, in PBS). After that, samples were incubated for 3 h at 37 °C in the dark. Samples were analyzed on a FACS-Calibur flow cytometer using the HTS module for processing 96-well plates. Cells were excited at 488 nM and fluorescence was collected with a FL2-A filter. The distributions of both FL2-A and FSC-H values were processed to find the two main density peaks, which correspond to the two cell populations in G1 and G2 phase of the cell cycle, as described in (Peter et al., 2018).

Whole chromosome size was estimated for MAL at 2 and 100 scb through Pulsed-Field Gel Electrophoresis, as described in (Louis, 1998).

### Generation of *MRE11* and *TBF1* mutants

The deletion of *MRE11* was performed by standard homologous recombination. The *URA3*-containing cassette was amplified from the laboratory S288C strain carrying a functional *URA3* gene. 80-bp primers used for the amplification were designed to contain the first and last 20 bp of *URA3* followed by 60 bp of homology with flanking regions upstream and downstream of *MRE11*. Samples were transformed using the standard lithium-acetate protocol, plated on selective medium lacking uracil and incubated at 30 °C for 3–7 days. Candidate transformed clones were validated by PCR using one primer designed inside *URA3* and another primer designed on the outside region of *MRE11*.

The deletion of one copy of *TBF1* in humanized strains carrying two chromosome XVI copies was performed by standard homologous recombination. The *HYGMX*-containing cassette was amplified from a plasmid carrying this antibiotic marker. 80-bp primers used for the amplification were designed to contain 20 bp of the MX region followed by 60 bp of homology with flanking regions upstream and downstream of *TBF1*. Samples were transformed using the standard lithium-acetate protocol, plated on selective medium containing hygromycin (200 ug/mL) and incubated at 30 °C for 3–7 days. Candidate transformed clones were validated by PCR using a combination of two PCR reactions: one reaction carried one primer designed inside *HYGMX* and another primer designed on the outside region of *TBF1*; the other reaction carried both primers designed inside *TBF1*. Only clones positive for both reactions were selected, as we wanted to delete only one copy of *TBF1*.

The introduction of *MRE11* Q447P, as well as the introduction of *TBF1* in the *HO* locus, were performed by using CRISPR/Cas9 genome editing. The plasmid harboring Cas9 was obtained from Addgene pUDP004 and linearized with BsaI. The resistance to acetamide was replaced with the resistance cassette to hygromycin which was amplified from a plasmid harboring the *HYGMX* gene. The gRNAs with the necessary nucleotides for self-cleavage were designed using the software UGENE and ordered as synthetic oligos from Eurofins Genomics (TM). Plasmid constructions were performed as described in (Mozzachiodi et al., 2022). The cassette used for the engineering of *MRE11* Q447P was designed to overlap the mutation site. The cutting site for Cas9 was modified to make it unrecognizable, by replacing one nucleotide with another one that did not change the aminoacid of the corresponding codon. The single-stranded parts of this cassette were ordered as a unique synthetic oligo at Eurofins Genomics. They were subsequently mixed at equimolar ratio, heated at 95 °C for 15 minutes and cooled down at room temperature to be used for the transformation. The cassette used for the knock-in of *TBF1* into the *HO* locus was amplified from the humanized ancestor at 0 scb (MJD548). 80-bp primers used for the amplification were designed to contain the first and last 20 bp of *TBF1* followed by 60 bp of homology with flanking regions upstream and downstream of *HO*. Yeast samples were transformed using the standard lithium-acetate protocol. Cells were plated on selective medium containing hygromycin (200 μg/mL) and incubated at 30 °C for 3–7 days. *MRE11* Q447P candidate transformed clones were validated by Sanger sequencing of selected colonies. *ho::TBF1* candidate transformed clones were validated by PCR using one primer designed inside *TBF1* and another primer designed on the outside region of *HO*. Positive clones were streaked on Leucin/Tryptophan dropout medium and grown for 2 days at 30 °C to allow plasmid loss and stored at -80 °C in 25% glycerol stocks.

### Quantitative PCR to track chrXVI copy number

To track the occurrence of chrXVI aneuploidy during MAL, we extracted DNA from yeast cells at various timepoints during the MAL protocol using a MasterPure Yeast DNA purification kit (Epicentre), following the manufacturer’s instructions. Primers were designed for the *TBF1* locus and for one control locus (*PWP2*) on chromosome III, since this chromosome never underwent aneuploidy in any line. To determine the copy number of chrXVI blocks in 8 single clones derived from Hum11 at 31 st (MJD460), we extracted DNA from the clones and designed one pair of primers for each block. Primers were designed on the following loci: *FUM1*-Block1; *TBF1*-Block2; *RPC40*-Block 3; *NUT2*-Block4. We also used the same control locus on chrIII (*PWP2*). DNA was diluted to 8 ng/uL for all the samples and the qPCR was performed using iTaq Universal SYBR Green Supermix (total volume: 10 µL) and run on an Applied Biosystems StepOne machine using the comparative ΔCt method. The PCR protocol was: initial denaturation (95 C; 15 min) followed by 45 cycles of: denaturation (95 C; 15 s), annealing (60 C; 30 s), extension (72 C; 30 s). We performed two replicates per sample per pair of primers. We calculated the copy number of the chrXVI regions covered by the primers by first subtracting the Ct of the chrIII control to all the primer pairs (ΔCt). We further normalized each primer pair’s ΔCt by subtracting the corresponding ΔCt of an internal control strain that has one copy of each region (humanized ancestor at 0 scb-MJD548). We calculated the copy number (CN) of each region as *CN=2^-ΔΔCt^*. In addition to the investigated samples, we also included a humanized line aneuploid for the full length of chromosome XVI (Hum3-MJD420) and the humanized ancestor at 0 scb (MJD548) that has one copy of each chromosome, as positive and negative controls, respectively.

### DNA extraction and short-read sequencing

Cells were inoculated in 5 ml of liquid Leucin/Tryptophan dropout medium and grown overnight at 30 °C in a shaking incubator. DNA was extracted using “Yeast Masterpure” kit (Epicentre, USA) following the manufacturer’s instructions. Illumina paired-end libraries (2×150 bp) were prepared according to manufacturer’s standard protocols and sequenced with an HiSeq 2500 instrument, at the NGS platform of Institut Curie. For the MAL experiment, two initial strains at 2 scb (one with the humanized background and one with the wild-type background) and all the strains at 100 scb were sequenced. For the AEL experiment, all the 5 humanized and 2 wild-type lines at 31 ST were sequenced. We additionally sequenced the humanized and wild-type ancestors at 0 scb, plus 8 single clones derived from Hum11 at 31 st. DNA from these 10 samples was extracted as described above but sequencing was performed by Novogene (Cambridge, UK).

### Long-read sequencing and structural variant analysis

Cells from a humanized line at 2 scb (Hum1), 4 humanized lines at 100 scb (Hum1,4,8,15) and two single clones derived from a AEL started from 100 scb (Hum11 at 31 st) were grown for two nights in liquid Leucin/Tryptophan dropout medium at 30 °C. Genomic DNA was extracted using “Yeast Masterpure’’ kit (Epicentre, USA) according to the manufacturer’s instructions. No vortexing was performed during the DNA extraction to avoid excessive DNA fragmentation. The correct size distribution of DNA fragments after the DNA extraction was verified by gel electrophoresis using the lambda DNA as molecular weight size marker. The MINION (Oxford Nanopore) sequencing library was prepared using the SQK-LSK109 sequencing kit according to the manufacturer’s instructions. The library was loaded onto a FLO-MIN106 flow cell and sequencing was run for 72 hours. Long read basecalling, demultiplexing and scaffolding were performed using the pipelines LRSDAY and LRSDAY_Patch (https://github.com/nicolo-tellini/LRSDAY-Patch) and the assembler was set as “canu” (Yue & Liti, 2018). Dotplots were generated using *nucmer* and *mummerplot* (Marçais et al., 2018). Structural variants were further confirmed by manually inspecting reads encompassing the breakpoints using IGV (v2.3.68) (Thorvaldsdottir et al., 2013).

### Short-read data preprocessing and CNVs estimation

We mapped Illumina reads resulting from MAL and AEL on the *S. cerevisiae* SGD reference genome assembly using *bwa mem* (v0.7.12) (Li & Durbin, 2009). Read mapping runs were stored in the Sequence Alignment/Map (SAM) format and its binary form (BAM) and post-processing steps were performed by picard tools (v2.8.0). Sorting, indexing and per-position coverage calculation were performed using SAMtools (v1.2) (Li et al., 2009). Regions with higher coverage values as compared to the genome-wide median were considered as copy number variants and their effective copy number was calculated as the ratio between their median coverage and the median genome-wide coverage for the strain in which they were observed. The ORFs copy number was estimated as in (Peter et al., 2018). Reads from each strain were mapped against the reference S288C genome from the SGD database using *bwa mem* (Li & Durbin, 2009), to obtain the baseline coverage. Then, the same reads were mapped against the pangenome file from (Peter et al., 2018) using *bwa mem* and the option *-U 0*, and the median coverage was extracted for each pangenomic ORF. Coverages from the bam files were extracted using a custom script using the R library *Rsamtools*.

We used the software freebayes (v0.9.5) to call markers from the BAM files. Results were stored in Variant Call Format (VCF) files. We filtered the VCF files to keep only SNVs/indels with quality higher than 20 and coverage higher than 10.

### Identification of *de novo* SNVs in MAL and AEL

For MAL, we subtracted the SNVs/indels in the humanized strain at 2 scb (MJD1) from the humanized lines at 100 scb, and the SNVs/indels in the wild-type strain at 2 scb from the wild-type lines at 100 scb, using the vcf-isec program included in the suite VCFtools (v0.1.16). For AEL, we subtracted the SNVs/indels in the corresponding mutation accumulation lines at 100 scb from their derived AEL. Next, we searched for shared SNVs present in at least two MAL at 100 scb (or in two AEL for the AEL experiment) and removed them from the VCF files, based on the assumption that identical *de novo* SNVs are highly unlikely to arise independently and derive from incorrect mapping or poor coverage. All the remaining SNVs/indels were manually validated using IGV (v2.3.68) (Thorvaldsdottir et al., 2013). SNVs were annotated using the *S. cerevisiae* SGD genome assembly implemented in the online software Variant Effect Predictor (VEP) (McLaren et al., 2016). Further annotation of the functional impact of SNVs was carried out using mutfunc (Wagih et al., 2018). We tested whether selection during MAL might have affected base substitution rates. The proportion of observed vs expected coding SNVs (74 *vs* 75 %) and non-synonymous SNVs (68 *vs.* 75 %) in wild-types supports a close-to-neutral mutation accumulation scenario (binomial test: *p*=ns), while the corresponding proportions in humanized yeasts (59 and 76 % for coding and non-synonymous mutations, respectively) indicate an effect of selection in this group, with mutations less likely to occur in coding regions (binomial test: *p*<0.0001). *De novo* variants derived from MAL were used to estimate base substitution and insertion/deletion rates (*μ*) as *μ*=*(n/bp)/g*, where *n* is the number of *de novo* SNVs present in each line, *bp* is the (haploid) genome size of the *S. cerevisiae* SGD genome assembly and *g* is the number of generations performed in the mutation accumulation experiment (2140 for wild-type lines and 1640 for humanized lines).

### Experimental validation of *de novo* SNVs in MAL and AEL

Two SNVs from humanized MAL were validated by Sanger sequencing. One variant was located inside the gene *MEC1* (Hum4: *MEC1* R1824H) and the other was inside the gene *TEL1* (Hum8: *TEL1* L2308R). A pair of primers (upstream and downstream) was designed for each SNV using Unipro UGENE (Okonechnikov et al., 2012), and used to perform Polymerase Chain Reaction (PCR). We checked the presence of the mutations at all timepoints during the MAL protocol. PCR products were sequenced by Eurofins Genomics. The presence and the genotype of the variants were checked by visual inspection of the electropherograms. We confirmed the presence of both variants and discovered that the mutations in *TEL1* occurred between 60 and 80 scb, while the mutation in *MEC1* occurred between 40 and 60 scb. We used the same procedure to confirm the presence of *MRE11* and *RAD50* mutations in AEL started from 100 scb (Hum1: *MRE11* K477I; Hum2: *MRE11* Q447P; Hum3: *RAD50* L171P).

### Analysis of *TLC1* variation

To analyze the extent of genetic variation in the *TLC1* template and its flanking regions across the *Saccharomyces sensu stricto* species and across the *S. cerevisiae* population, we first downloaded published *TLC1* sequences from (Liti et al., 2006), spanning several *S. cerevisiae* and *S. paradoxus* isolates. We then downloaded the *TLC1* sequence from the *Saccharomyces* Genome Database (SGD) and used it to search for orthologs in other *Saccharomyces sensu stricto* species in the NCBI database, using the *blastn* algorithm. We then performed a multiple sequence alignment using *clustal omega* (https://www.ebi.ac.uk/jdispatcher/msa/clustalo). Next, we gathered publicly available Illumina sequencing datasets relative to 3,139 *S. cerevisiae* strains, and further sequenced 241 new *S. cerevisiae* isolates (unpublished data). The latter were sequenced at The Earlham genomics facilities (www.earlham.ac.uk) according to the LITE pipeline (Perez-Sepulveda et al., 2021). We run whole-genome short-reads alignments with *bwa mem* (v.0.7.17-r1198-dirty) (Li, 2013) and processed the bam files according to the latest samtools guidelines (samtools fixmate, sort, markdup and view) (v1.19). For each isolate, we produced a gVCF file using bcftools (mpileup, call, view and annotate) (v1.19) and extracted the genotypes for the 16 nt template (S288C v.R64-3-1_20210421, chrII:308054-308069) and 20 nt upstream and downstream this sequence (chrII:308034-308053, chrII:308070-308089). We converted the gVCF genotypes in fasta sequences concatenating the alleles across consecutive positions with vcf2phylip.py (v.2.8) and created a multifasta file. The heterozygous positions were resolved according to the IUPAC nomenclature. At the end, we extracted the unique sequences, their occurrences and the list of isolates sharing the unique sequences with seqkit rmdup (v.2.4.0) (W. Shen et al., 2016). The output was parsed with custom R scripts.

### Telomere length, ITS content and Y’ copy number estimation from short-read sequencing

Telomere length, ITS content and copy number of Y’ elements were determined using Y^ea^ISTY (D’Angiolo et al., 2023). Telomere length and ITS content were measured separately for the TG_1-3_ and T_2_AG_3_ motifs. For TG_1-3_ we used the published version of Y^ea^ISTY, while we developed a new version compatible with the T_2_AG_3_ motif for this study. We used a threshold of 20 bp of telomeric repeats stretch length for TG_1-3_, and a threshold of 18 bp for T_2_AG_3_. By comparing our results with the estimates derived from long-read genome assemblies, we determined that our lengths were underestimated and the underestimation was particularly pronounced for the T_2_AG_3_ repeat. This is likely due to the fact that the T_2_AG_3_ motif is longer and more complex than TG_1-3_ and sequencing errors are more prone to disrupt the continuity of T_2_AG_3_ stretches. As a consequence, we avoid direct comparison of total telomere lengths (TG_1-3_+T_2_AG_3_) between humanized and wild-type yeasts.

### Chromosome-ends annotations from long-read genome assemblies

X and Y’ elements were annotated using LRSDAY (Yue & Liti, 2018). Elements annotated as “partial” were further inspected by extracting the corresponding and surrounding genomic regions (∼10 kb) with *bedtools getfasta* and by performing a *blastn* local alignment with the SGD reference genome. Most of the partial elements were instead full versions and were re-annotated using the coordinates derived from the local alignment. We annotated TG_1-3_ and T_2_AG_3_ repeats in the genome assemblies by using custom Perl scripts. We merged the repeats which were interspaced by only 1 bp into a unique stretch by using the software mergeBed included in the suite bedtools (version 2.17.0) with the option “-d 1” (Quinlan & Hall, 2010). Then, we filtered the list to keep only the stretches longer than 10 bp. The interstitial stretches were annotated as ITS if they closely preceded a Y’ element, while the stretches at the termini were annotated as telomeres. We further mapped long reads derived from each strain onto its corresponding genome assembly using *bwa mem* (v0.7.12) and manually visualized long reads encompassing telomeres using IGV (v2.3.68) (Thorvaldsdottir et al., 2013).

### Mass spectrometry and proteomics data analysis

*Cultivation of library:* strains were randomized manually in triplicates onto 96-well plates, with the exception of strains MJD436-447, MJD450-455, MJD481-500, and MJD586-597, each of which was cultivated once. Strains were pinned onto solid Leucine/Tryptophan dropout agar using a Singer Rotor and incubated at 30 °C for 48 h. Subsequently, each colony was pinned into 200 µL of Leucin/Tryptophan liquid dropout medium and the samples were incubated at 30 °C without shaking. After 32 h, these pre-cultures were mixed thoroughly by pipetting and 80 µL of each sample were added to 1.55 mL of pre-warmed Leucin/Tryptophan liquid dropout medium in 96-well deep-well plates. One sterilized borosilicate glass bead was added per well in order to aid mixing. Plates were sealed with a Breathe-Easy membrane and incubated for 14.5 h at 800 rpm, 30 °C. Then, 1.4 mL of culture were transferred to a new deep-well plate, and cells were harvested by centrifugation (2880 xg, 5 min, 4 °C). The optical density at harvest was measured using an Infinite M Nano plate reader (Tecan). The supernatant after centrifugation was removed by inverting the plate, and cell pellets were immediately cooled on dry ice. Cell pellets were stored at -80 °C until lysis.

*Proteomic sample preparation*: samples for proteomics were prepared in a 96-well high-throughput format as previously described (Muenzner et al., 2024). Briefly, cells were lysed in 200 µL 100 mM ammonium bicarbonate, 7 M urea by beat beating using a Geno/Grinder (Spex). Subsequently, samples were reduced and alkylated using DTT (20 µL, 55 mM) and iodoacetamide (20 μL, 120 mM), respectively. Samples were then diluted by adding 460 µL of mL 100 mM ammonium bicarbonate. 500 µL per sample were digested using 2 µg Trypsin/LysC (Promega, Cat#V5072). After 17 h of incubation at 37 °C, 25 µL 20% formic acid were added to the samples, and peptides were purified using solid-phase extraction as described before (Messner et al., 2023). Eluted samples were vacuum-dried and subsequently reconstituted in 90 µL 0.1% formic acid. An equivoluminal pool of all samples (5 µL per sample) was generated to be used as technical controls during MS measurements. Peptide concentrations of samples and the pool were determined using a fluorimetric peptide assay kit (Thermo Scientific, Cat#23290). Samples with a too low optical density at harvest (OD600 < 0.25) and/or too low peptide concentrations (no peptide signal observed) were excluded from mass spectrometric measurements.

*LC–MS/MS measurements*: samples were analyzed in two batches (96 samples per batch plus analytical quality control (pool) injected every ten samples) on a SCIEX ZenoTOF 7600 System mass spectrometer coupled to a Waters ACQUITY UPLC M-Class System. Prior to MS analysis, 200 ng peptides were chromatographically separated with a 60 min gradient on a Waters HSS T3 column (300um x 150mm, 1.8um) heated to 35 °C, using a flow rate of 5 µL/min where mobile phase A and B were 0.1% formic acid in water and 0.1% formic acid in acetonitrile, respectively. For data independent acquisition, a zenoSWATH MS/MS acquisition scheme with 85 variable size windows and 11 ms accumulation time was used. Ion source parameters were set to: ion source gas 1 & 2 – 12 and 60 psi respectively; curtain gas 25, CAD gas 7 and source temperature at 150°C; spray voltage – 4500V.

*Data pre-processing*: raw data were processed with DIA-NN (version 1.8, available from https://github.com/vdemichev/DiaNN) (Demichev et al., 2020) using the following settings: fragment ion m/z range set from 300-1800, mass accuracy set to 20 ppm at the MS2 and MS1 level, respectively, scan window set to 6, MBR (Match-between-runs) enabled, and Quantification strategy set as “Robust LC (high Precision)”. A spectral library-free approach and yeast reference proteome (UniProt UP000002311, accessed on 23.04.2021) were used for annotation. The resulting report.tsv file was further pre-processed at the precursor level in R. First, all non-proteotypic peptides were excluded and precursors with q.values, PG.q.values, Global.q.values, Global.PG.q.values, or GG.q.values > 0.01 were excluded. Next, samples with very low precursor numbers (Z-score < -2.5) were removed. Batch correction was performed by equalizing the median precursor quantity of each batch. Protein quantities were determined for all proteins with two or more precursors mapped using the diann_maxlfq() function of the DIA-NN R package (https://github.com/vdemichev/diann-rpackage), resulting in a dataset covering 3503 proteins across 181 samples. No imputation was applied to the proteomic dataset and undetected proteins were treated as missing values.

*Data analysis*: after these pre-processing steps, we calculated the median of the three technical replicates and normalized them using the wild-type strain 1 at 2 scb (MJD5) as a reference, to quantify the relative changes. The quality of the technical replicates was confirmed by the high Pearson’s correlation coefficients (>0.98) between each pair. To identify genes that are differentially expressed as compared to the reference, we calculated the fold-changes and the p-values corresponding to each gene using the edgeR library in R. The normalization factors for differences in measurement depth across different samples were calculated using the function ‘calcNormFactors’. Dispersion factors were estimated using the functions ‘estimateCommonDisp’, ‘estimateTagwiseDisp’ and ‘estimateTrendedDisp’ and we quantified the dispersion of the gene expression based on the mean values. The function ‘exactTest’ was used to identify significant differential expressions, which were then visualized as volcano plots highlighting significant fold-changes in expression using the ‘EnhancedVolcano’ function from the ‘EnhancedVolcano’ library. The Principal Component Analysis was performed using the function ‘fviz_pca_ind’ from the library ‘factoextra’ in R.

Next, we derived lists of differentially abundant proteins for each strain by selecting those showing a log_2_FC>=1 or log_2_FC<=-1 and *p*<=0.05. Each list was further filtered on the basis of the aneuploidies carried by each strain, in order to retain only proteins that are differentially abundant due to upregulated/downregulated expression and not by copy number modifications of their encoding genes located on duplicated chromosomes. Once we produced a list for each strain, we derived lists of differentially abundant proteins for humanized and wild-type yeasts by group (MAL at 40 and 100 scb, AEL from 2 and 100 scb) by selecting those present in >40% of the strains in each group and combined them, deriving 110 proteins whose abundance changed in at least one group as compared to the reference (**Fig. 3d**). In contrast, the list of proteins defining the THPR was derived by combining the proteins that are differentially abundant in at least 2 humanized MAL at 40 or 100 scb. The THPR was compared with a set of genes that are differentially expressed upon *cdc13* temperature-sensitive mutation and were downloaded from (Greenall et al., 2008), as well as with a list of Rap1p gene targets. The list of Rap1p targets was downloaded from SGD and derived from multiple studies (Brindle et al., 1990; Duveau et al., 2021; Hu et al., 2007; Lickwar et al., 2012; MacIsaac et al., 2006; McNeil et al., 1990; Platt et al., 2013, Song et al., 2020, Stanway et al., 1989).

### GO term analysis

Standard GO term analysis was performed separately on the variants extracted from wild-type and humanized MAL. Due to the small number of *de novo* variants occurred during AEL, we combined the variants in AEL from 2 and 100 scb and performed GO term analysis separately on the variants extracted from wild-type and humanized AEL. GO term analysis for *de novo* mutations was performed using the GO Term Finder tool available at *Saccharomyces* Genome Database (SGD). Significant GO terms were extracted by the algorithm implemented in the tool, with a False Discovery Rate (FDR) corrected α threshold of 0.1. GO term analysis for differentially abundant proteins was performed using the *gprofiler2* package in Rstudio (Kolberg et al., 2020).

