## Supplementary_Text_and_figures for "The adaptive molecular landscape of reprogrammed telomeric sequences"

### Supplementary discussion

#### 1, Structural variation in humanized MAL

We performed Pulsed Field Gel Electrophoresis (PFGE) on one representative wild-type and humanized line at 2 scb and all the MA lines at 100 scb to investigate how genome instability affected chromosome size. The lines at 2 scb and the wild-types at 100 scb showed the same band pattern, while many band shifts were evident in the humanized lines at 100 scb, indicating the occurrence of abundant structural variation during MAL. By comparing our Nanopore-derived ITS/Y' annotations and band positions we could recapitulate major chromosomal size increases compatible with the massive amplification of ITS/Y' elements, which is especially pronounced on chromosomes VIII and XVI (**Supplementary Figure 2d**). The simplest explanation for this trend is that chromosome-ends containing multiple Y' elements are preferentially amplified as the original Y' elements constitute a seed for tandem duplication. Y' elements in other chromosome ends that did not originally contain them may have arisen through recombinational spread, as previously observed (Liti et al., 2005).

We further investigated one humanized MA line that showed massive band shifts by applying long-read sequencing technology (Hum8 at 100 scb). The genome assembly revealed a chromosomal rearrangement involving chromosomes V and XV. Specifically, the terminal part of chromosome V-R (~130 kb) is attached to the inverted left part of chromosome XV (~560 kb). This rearrangement resulted in the internalization of chrV-R telomeres as ITS and it is stabilized by chrXV-L telomeres and its centromere (CEN15) (**Supplementary Figure 6a-b**). Pulsed-field gel electrophoresis and manual inspection of long reads encompassing the breakpoints confirmed this rearrangement and revealed that it was mediated by the *ADE2* gene, which is not only present at its native locus on chrXV (564476..566191) but it has also been introduced in a subtelomeric site on chrV-R to be used as a marker of telomeric silencing. It is unclear whether the remaining parts of chromosomes V and XV are attached to each other or constitute two independent chromosomes. Although we did not detect any long reads supporting this structure, the first scenario is more likely as the resulting recombinant chromosome would be stabilized by chromosome V centromere (CEN5) plus chrV-L and chrXV-R telomeres, whereas the second scenario would involve the *de novo* formation of a centromere and telomeres. The diploid nature of humanized line 8 makes it difficult to precisely disentangle its chromosomal structure, as duplicated sequences are notoriously difficult to assemble, and we cannot exclude the possibility that the two subgenomes have different chromosomal structures.

#### 2, The proteomic landscape of humanized yeasts across experimental evolution

We investigated the impact of telomere variation on the proteomic landscape of humanized and wild-type yeasts through analysis of protein abundance by mass spectrometry, using the wild-type ancestor at 2 scb (MJD5) as reference for normalization. The proteomic profiles of both the ancestors (0 and 2 scb) and the wild-type lines (MAL and AEL from 100 scb) were similar to the reference, whereas the humanized lines showed a higher number of proteins whose abundance changed by at least 2-fold ($log_2FC>1$ or $log_2FC<-1$, $p<0.05$). During MAL, humanized lines had an average of 147 and 77 differentially abundant proteins at 40 and 100 scb, respectively. Part of these returned to wild-type levels after AEL, while others remained differentially expressed. To gain a better insight on the mechanisms driving the "telomere humanization proteomic response" (THPR), defined as the proteins that are differentially abundant in at least 2 humanized MAL at 40 or 100 scb, we compared it with other datasets. First, we compared the THPR with a set of genes that are differentially expressed, at the transcription level, upon *cdc13* temperature-sensitive mutation (*n*=232) (Greenall et al., 2008). We observed a statistically significant overlap of 79 genes between the two datasets: 15 overlapped between the datasets in the same direction, while 64 overlapped in opposite directions (two-tailed $X^2$ test, $p$=6.5e$^{-20}$, **Tables S11-12**). Overlapping genes underlied stress response functions like dNTP synthesis (*RNR2*, *RNR4*), response to oxidative stress, trehalose metabolism and energy production. However, while the upregulation of dNTP synthesis genes is conserved between the two datasets, other stress response genes are regulated in opposite directions, being upregulated in the *cdc13* temperature-sensitive mutation dataset and downregulated in the THPR. The THPR also had some unique genes: the downregulated genes underlied functions in protein and RNA transport, membrane organization and aminoacid biosynthesis, whereas the upregulated ones underlied nucleotide biosynthesis functions.

We further investigated whether the THPR was enriched in proteins regulating telomere length by intersecting it with a list of reported Telomere-Length-Maintenance (TLM) genes present in our proteomics dataset (*n*=147). This analysis yielded 15 genes that are in common between the two lists. 11 genes were downregulated in the THPR (*PMR1, TPD3, SSH1, STO1, CTR9, MCD4, SEC63, XRN1, RAT1, HSP104, SAC1*) and 4 were upregulated (*RFA2, TSA1, BUD21, GCV3*). However, this overlap is not significantly higher than expected (two-tailed $X^2$ test, $p$=0.25) and these genes did not show any functional enrichment.

Upon telomere humanization, Rap1p is replaced by Tbf1p. This reconfiguration liberates Rap1p molecules that are therefore free to exert their additional role of transcription factors. We investigated whether the THPR might be mediated by the action of Rap1p as a transcription factor by intersecting the THPR with Rap1p targets (*n*=345). This analysis yielded 56 genes that are in common between the two lists (37 in the downregulated THPR and 19 in the upregulated THPR), although this overlap is not significantly higher than expected (two-tailed $X^2$ test, $p$=0.15). These genes underlied functions related to carbohydrate and nucleotide metabolism. Overall, these results show that the THPR is in part similar to the expression program triggered by telomere dysfunction, in that both programs trigger the upregulation of DDR target genes, but it also has its own specific signature (**Figure 3d and Tables S11-12**).

**3, Dosage compensation for genes located on duplicated chromosomes**

We compared the chromosome copy number inferred by short-read sequencing coverage and protein abundance Fold Change (FC) in a subset of strains for whom we possessed both types of data ($n$=44, excluding single clones derived from AEL of Hum11 that showed segmental duplications on chrXVI). While we observed a positive correlation between the two datasets across the 16 chromosomes ($r$=0.82, $p$<2.2e-16), there was a partial gene dosage compensation at the proteome level, with an average ratio of 1.92 between the coverage of aneuploid chromosomes and the genome-wide coverage, and an average FC of 1.59 between the proteome abundance of aneuploid chromosomes and that of the euploid reference strain (**Figure 5f**). Furthermore, we identified 3 humanized MAL at 100 scb (Hum3,4,7-chrXVI) and one humanized AEL (Hum1-chrXI) exhibiting mismatches between the genome and proteome copy number annotations, involving the loss of chrXI/XVI aneuploidies likely caused by an unstable karyotype.

We further investigated whether the aneuploidies observed in our MAL (chrXI, chrXVI) triggered a specific expression program by comparing the lists of differentially abundant proteins in euploid *vs* aneuploid lines for the two chromosomes. This analysis yielded two signatures of 45 and 29 proteins that are differentially abundant only in chrXI- and chrXVI-aneuploid lines, respectively. The chrXVI-aneuploid signature did not show any enrichment in specific functions, further confirming that the adaptive value of this aneuploidy lies in the protective function of Tbf1p towards the telomeres, and not in the triggering of a specific expression program. In contrast, the chrXI-aneuploid signature was enriched in stress response functions, suggesting that it may also have an adaptive value linked to its effect on stress response genes.

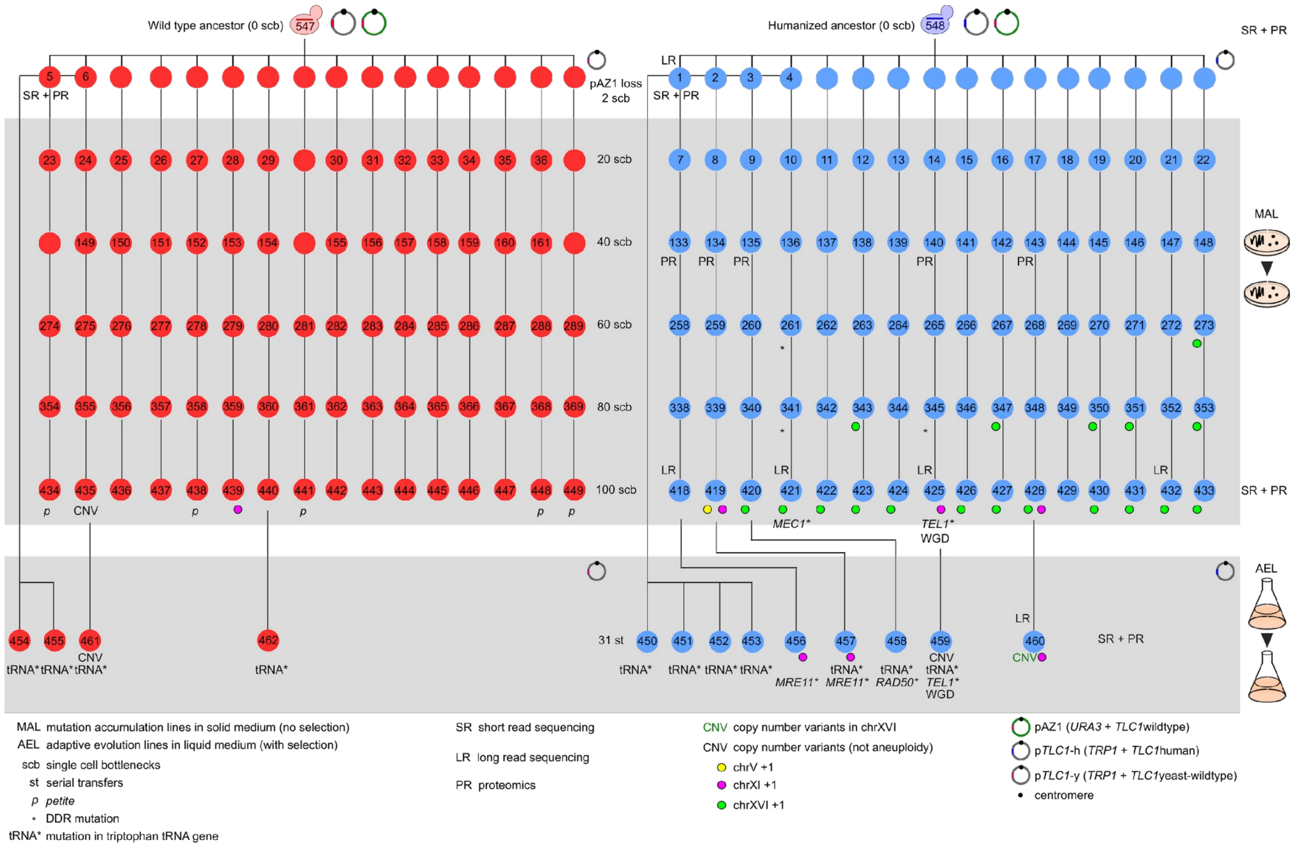

**Supplementary Figure 1 – Experimental evolution roadmap.** Detailed representation of experimental protocols used to evolve wild-type (red) and humanized (blue) ancestor strains in the absence (MAL-top panel) or in the presence (AEL-bottom panel) of selection. Symbols juxtaposed to single strains indicate main genetic/genomic features characterizing that strain or technologies (sequencing and/or proteomics) that have been applied to it. Numbers inside the circles indicate the strain ID as listed in Table S1. Humanized and wild-type lines are numbered from left (1) to right (16) and their derivation is indicated by plain vertical lines connecting the circles. Wild-type lines that became *petite* during MAL (indicated with *p*) were not further studied. The numbering of the lines is maintained from the MAL to the AEL protocol.

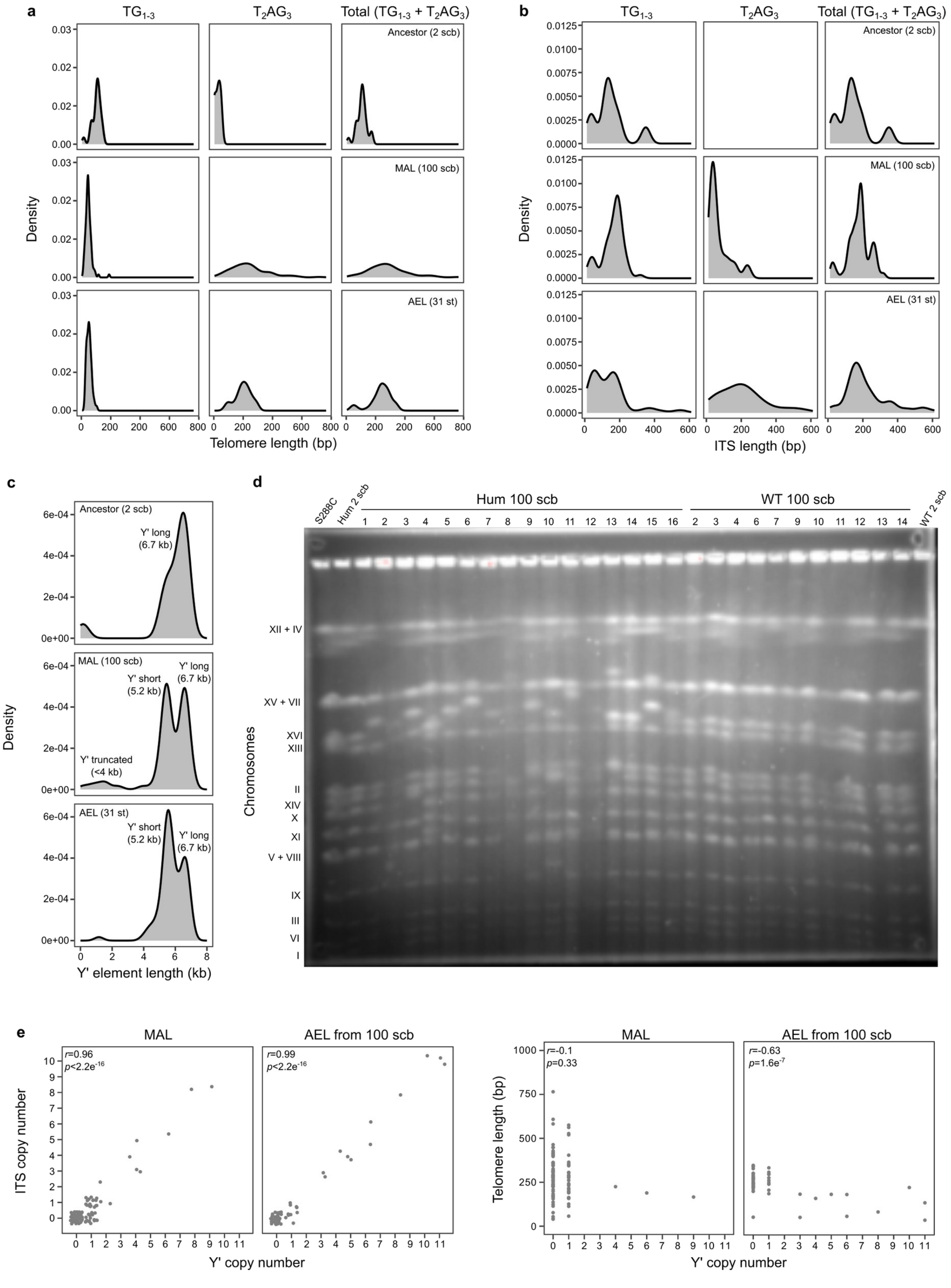

**Supplementary Figure 2 – Telomere, ITS and Y' length variation. a-b,** Size density plots of telomere and ITS length in Nanopore-sequenced lines, shown separately for $TG_{1-3}$, $T_2AG_3$ and total $TG_{1-3}$+$T_2AG_3$. $TG_{1-3}$ and $T_2AG_3$ segments in mixed, sandwich and double sandwich ITS are counted separately. For clarity, two telomeres with extremely long size (>1000 bp) are not shown in **a**. **c**, Size density plots of Y' length in Nanopore-sequenced lines. Peaks corresponding to long-version (6.7 kb) and short-version Y' (5.2 kb) are highlighted. Y' shorter than 4 kb are only present at truncated chromosome ends and derive from incomplete annotation. **d**, PFGE image of wild-type and humanized lines at the beginning (2 scb) and at the end (100 scb) of the MAL protocol. Lanes contain, from left to right: 1) a S288C laboratory control strain, 2) a humanized line at 2 scb, 3-18) humanized lines at 100 scb, 19-29) wild-type lines at 100 scb, 30) a wild-type line at 2 scb. The band patterns of the S288C control, WT 2 scb, Hum 2 scb and wild-types at 100 scb are identical, while the humanized lines at 100 scb have multiple band shifts involving mostly the chromosome XVI band, which increased its molecular weight in the majority of the humanized lines. There is extensive correlation between band shifts and ITS/Y’ amplifications shown in Figure 2e (e.g. Hum4 chrVIII). The humanized line 8, carrying a loss-of-function mutation in *TEL1*, shows an extensive band pattern variation with respect to the rest, compatible with multiple chromosomal rearrangements and subtelomeric amplifications. **e**, Correlation between ITS, Y' copy number and telomere length across Nanopore-sequenced MAL and AEL. The ancestor at 2 scb is not included in this analysis. Telomere length is calculated as the sum of $TG_{1-3}$ and $T_2AG_3$ repeats.

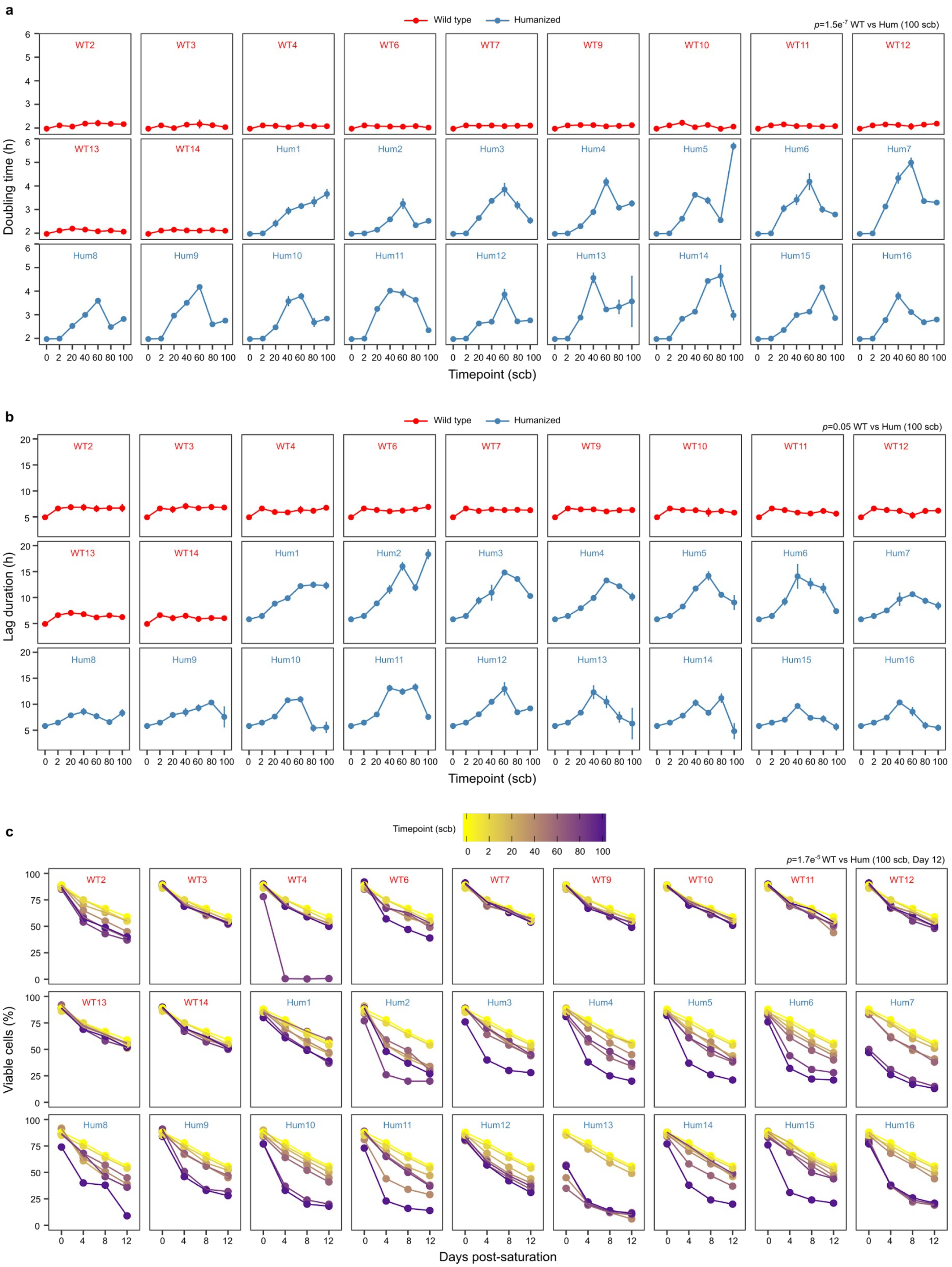

**Supplementary Figure 3 – Fitness dynamics in single MA lines. a-c**, Population doubling time (top), lag duration (middle) and survival rate (bottom) of single wild-type (red) and humanized (blue) MAL every 20 scb. Doubling time and lag measurements were performed in triplicate. The points represent average +- standard deviation among the replicates. Survival rate measurements were performed every 4 days in one replicate and the color tone (from yellow to purple) indicates the advancement in the MAL protocol (scb). Indicated *p*-values result from the comparison of combined wild-type and humanized lines at 100 scb (two-tailed Wilcoxon test).

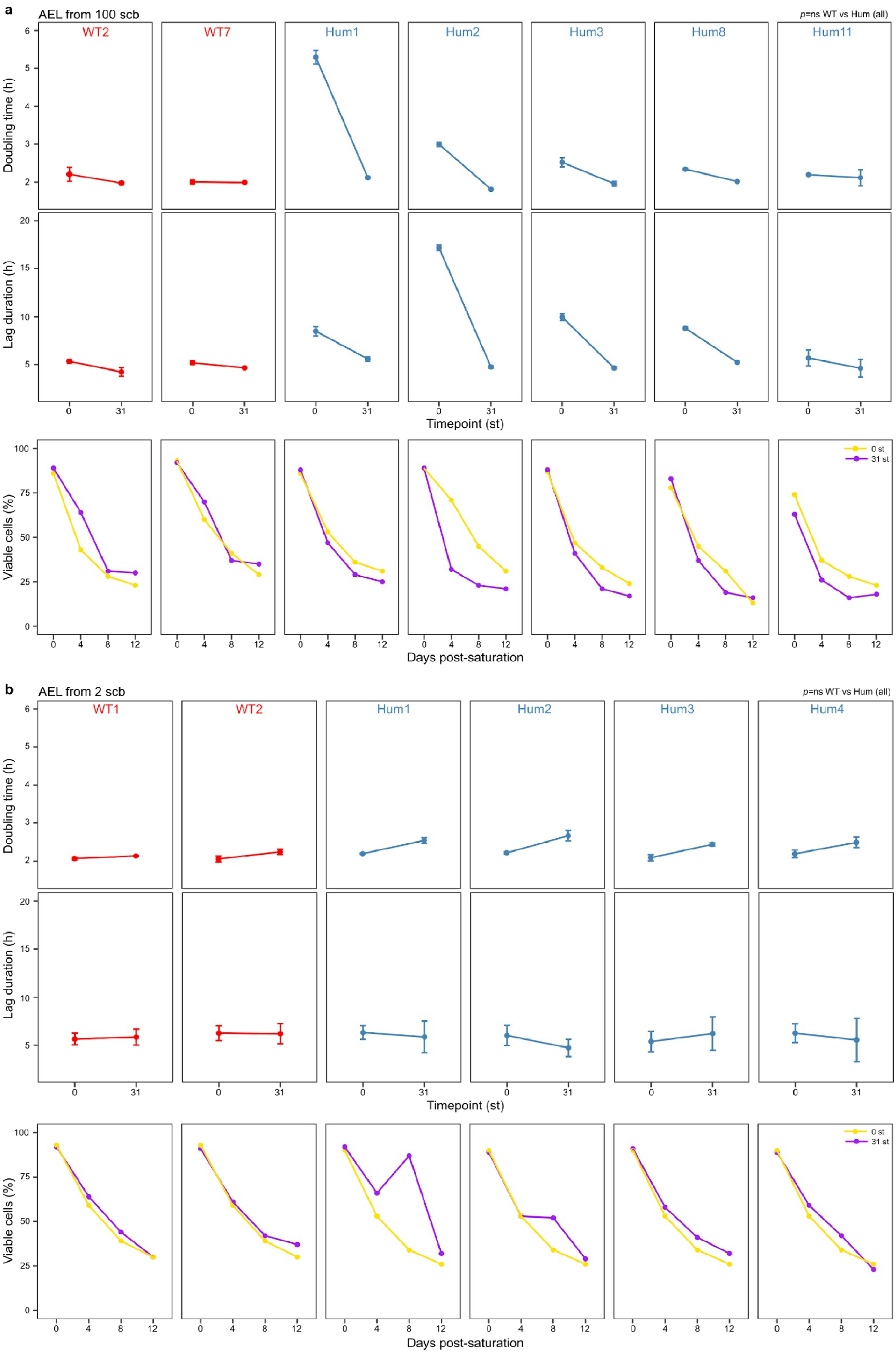

a
AEL from 100 scb
p=ns WT vs Hum (all)
WT2
WT7
Hum1
Hum2
Hum3
Hum8
Hum11
Doubling time (h)
Lag duration (h)
Timepoint (st)
Viable cells (%)
0 st
31 st
Days post-saturation
b
AEL from 2 scb
p=ns WT vs Hum (all)
WT1
WT2
Hum1
Hum2
Hum3
Hum4
Doubling time (h)
Lag duration (h)
Timepoint (st)
Viable cells (%)
0 st
31 st
Days post-saturation

**Supplementary Figure 4 – Fitness dynamics in single AE lines. a-c**, Population doubling time (top), lag duration (middle) and survival rate (bottom) of single wild-type (red) and humanized (blue) lines before (0 st) and after (31 st) AEL. Doubling time and lag measurements were performed in triplicate. The points represent average +- standard deviation among the replicates. Survival rate measurements were performed every 4 days in one replicate and the color indicates the advancement in the AEL protocol (st). Indicated *p*-values result from the comparison of combined wild-type and humanized lines at 31 st (two-tailed Wilcoxon test).

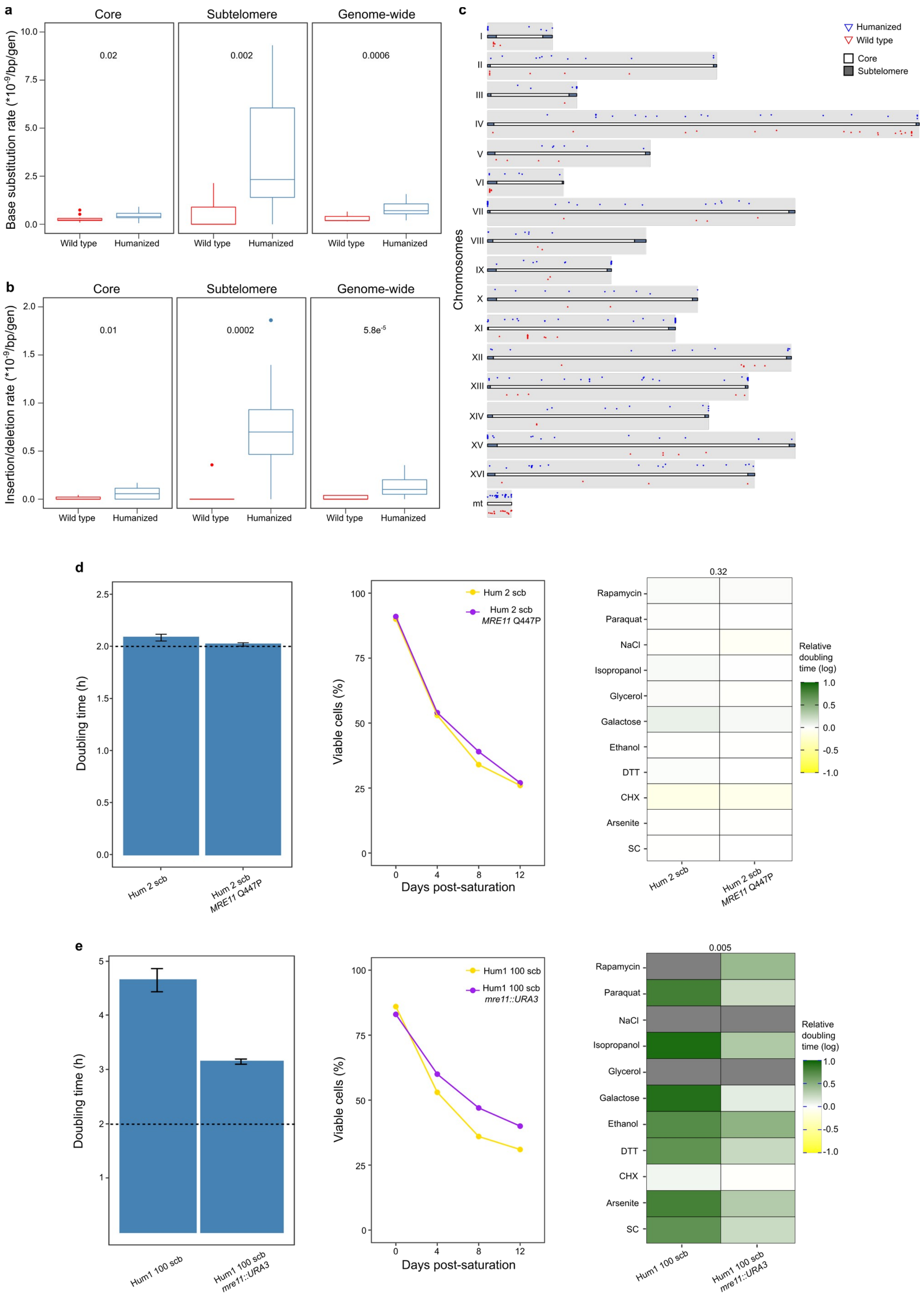

**Supplementary Figure 5 – The effect of DDR mutations on fitness. a-b**, Base substitution (**a**) and insertion/deletion rate (**b**) per bp per generation in wild-type and humanized MAL (n=11 and n=16,respectively). Boxplots as in Fig. 2. Numbers in the plots represent p-values resulting from the comparison ofwild-type and humanized MAL at 100 scb (two-tailed Wilcoxon test). **c**, Genome-wide distribution of SNVs and insertions/deletions in the humanized (upper line, blue spots) and wild-type (bottom line, red spots) MAL. Genome coordinates are based on the *S. cerevisiae* SGD genome reference. Dark grey areas at the chromosome tips represent subtelomeres and their boundaries are based on (Yue et al., 2017). The presence of multiple spots on the same vertical line denotes a high density of mutations in a restricted area, and it is especially frequent in subtelomeres. **d**, Population doubling time, survival rate and relative doubling time of a humanized ancestor at 2 scb (Hum1) before and after the introduction of a missense mutation in *MRE11* (Q447P). The dashed line indicates the population doubling time of wild-type. Color codes and number of replicates are as described in Figure 3. Population doubling time measurements were performed in triplicate. Survival rate measurements were performed in one replicate. Relative doubling time measurements were performed in 24 replicates for SC and 12 replicates for all the other conditions. Barplot: mean + standard deviation. **e**, Population doubling time, survival rate and relative doubling time of one humanized line at 100 scb before and after the replacement of *MRE11* by *URA3*. The dashed line indicates the population doubling time of wild-type. Color codes and number of replicates are as described in Figure 3. Barplot: mean + standard deviation.

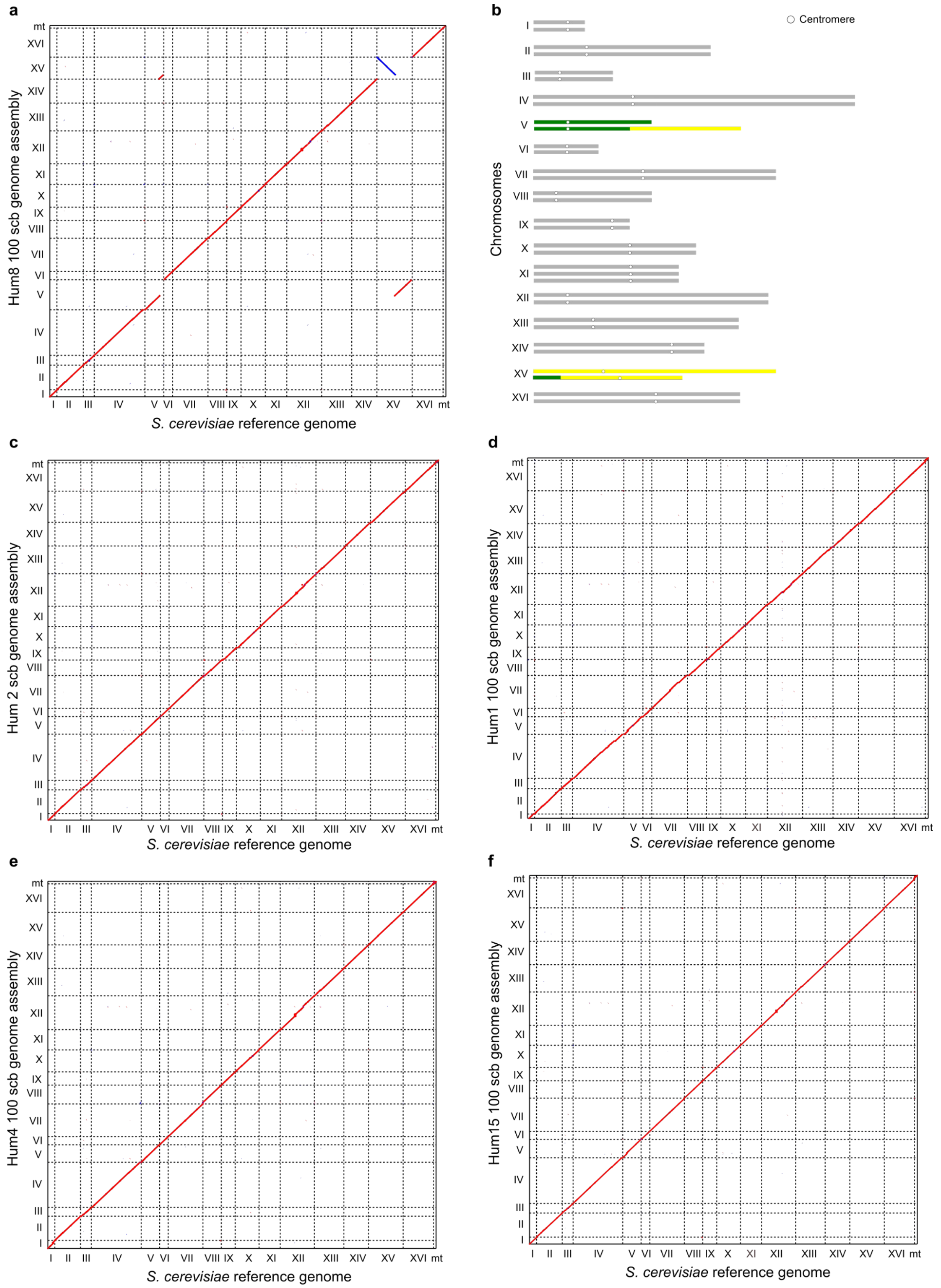

**Supplementary Figure 6 – Genomic structure of humanized MAL. a-f**, Comparison of the *S. cerevisiae* reference genome (x axis) and our Nanopore-derived genome assemblies (y axis) for a humanized ancestor at 2 scb (Hum1) and 4 MAL at 100 scb (Hum1,4,8,15). Sequence homology signals are indicated in red (forward match) or blue (reverse match). Black arrows indicate breakpoints of chromosomal rearrangements in the humanized line 8. Plot **b** represents a putative karyotype of humanized line 8 based on the genome assembly shown in **a** and the manual inspection of Nanopore reads encompassing the breakpoints of the rearrangements. White circles denote centromeres and colored chromosomes (V-green and XV-yellow) are the ones involved in rearrangements.

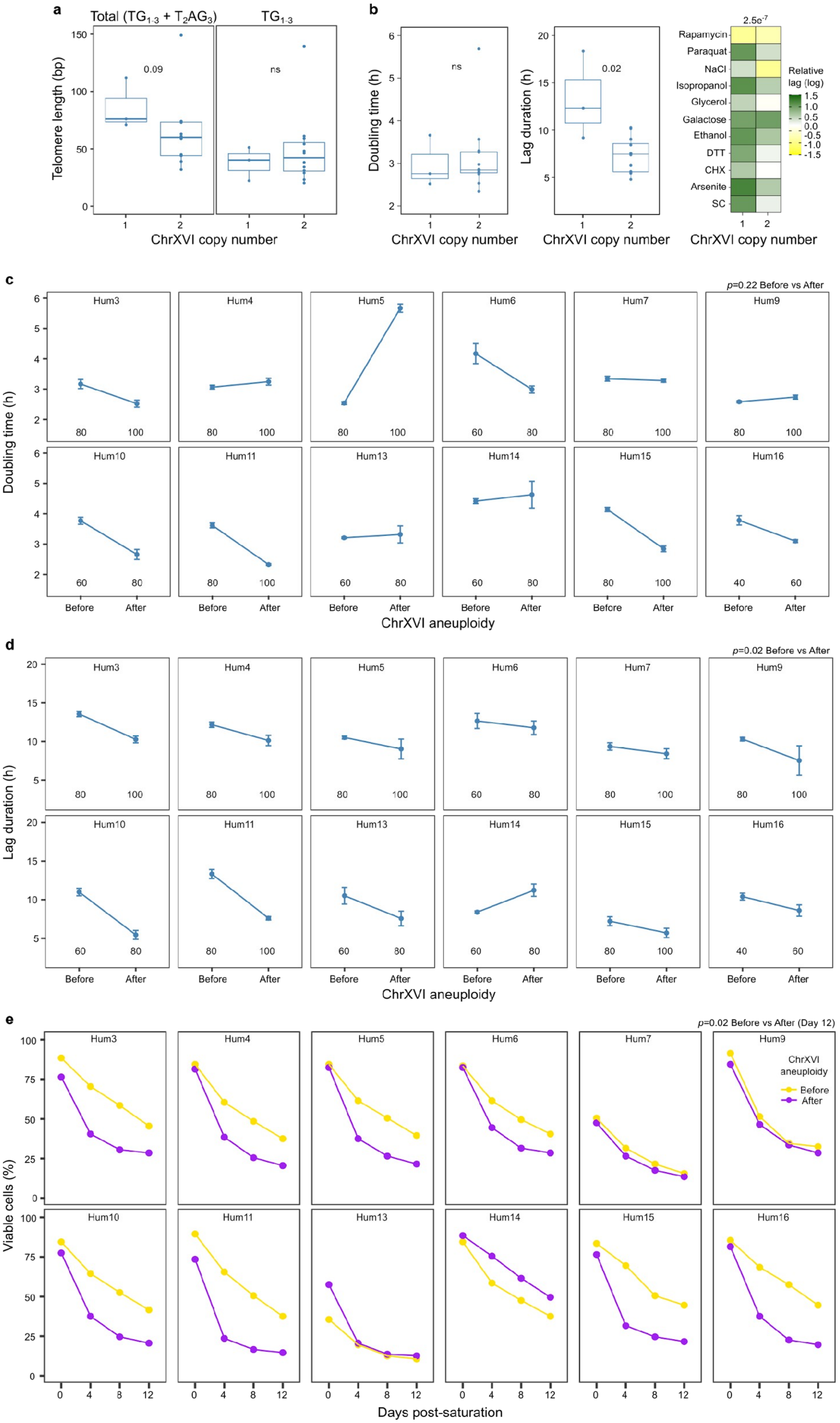
a
Total (TG1-3 + T2AG3)
TG1-3
Telomere length (bp)
150
100
50
0
0.09
ns
1
2
1
2
ChrXVI copy number
b
Doubling time (h)
6
5
4
3
2
ns
1
2
ChrXVI copy number
Lag duration (h)
20
15
10
5
0.02
1
2
ChrXVI copy number
2.5e-7
Rapamycin
Paraquat
NaCl
Isopropanol
Glycerol
Galactose
Ethanol
DTT
CHX
Arsenite
SC
Relative lag (log)
1.5
1.0
0.5
0.0
-0.5
-1.0
-1.5
1
2
ChrXVI copy number
c
p=0.22 Before vs After
Doubling time (h)
Hum3
Hum4
Hum5
Hum6
Hum7
Hum9
Hum10
Hum11
Hum13
Hum14
Hum15
Hum16
Before
After
ChrXVI aneuploidy
d
p=0.02 Before vs After
Lag duration (h)
Hum3
Hum4
Hum5
Hum6
Hum7
Hum9
Hum10
Hum11
Hum13
Hum14
Hum15
Hum16
Before
After
ChrXVI aneuploidy
e
p=0.02 Before vs After (Day 12)
Viable cells (%)
Hum3
Hum4
Hum5
Hum6
Hum7
Hum9
ChrXVI aneuploidy
Before
After
Hum10
Hum11
Hum13
Hum14
Hum15
Hum16
0
4
8
12
Days post-saturation

**Supplementary Figure 7 – Fitness dynamics upon chromosome XVI aneuploidy. a,** Telomere length proxy (total and $TG_{1-3}$-only) in humanized MAL that are either euploid (one copy) or aneuploid (two copies) for chromosome XVI. Boxplots as in Fig. 2. Color codes and number of replicates are as described in Figure 3. Humanized line 8 has been excluded from this analysis due to its diploidy. Numbers in the plots represent *p*-values resulting from the comparison of humanized lines from the underlying genetic backgrounds (two-tailed Wilcoxon test). **b,** Population doubling time (left), lag duration (middle) and relative lag (right) in humanized MAL that are either euploid (one copy) or aneuploid (two copies) for chromosome XVI. Boxplots as in Fig. 2. Color codes and number of replicates are as described in Figure 3. Humanized line 8 has been excluded from this analysis due to its diploidy. Numbers in the plots represent *p*-values resulting from the comparison of humanized lines from the underlying genetic backgrounds (two-tailed Wilcoxon test). **c-e**, Population doubling time (top), lag duration (middle) and survival rate (bottom) of single humanized lines before and after the duplication of chromosome XVI. Numbers inside the boxes indicate the timepoints corresponding to the change in chrXVI copy number. Doubling time and lag measurements were performed in triplicate. The points represent average +- standard deviation among the replicates. Survival rate measurements were performed every 4 days in one replicate and the color indicates the timing respect to the occurrence of the aneuploidy (before/after). Indicated *p*-values result from the comparison of combined humanized lines before and after chrXVI aneuploidy (two-tailed Wilcoxon test).

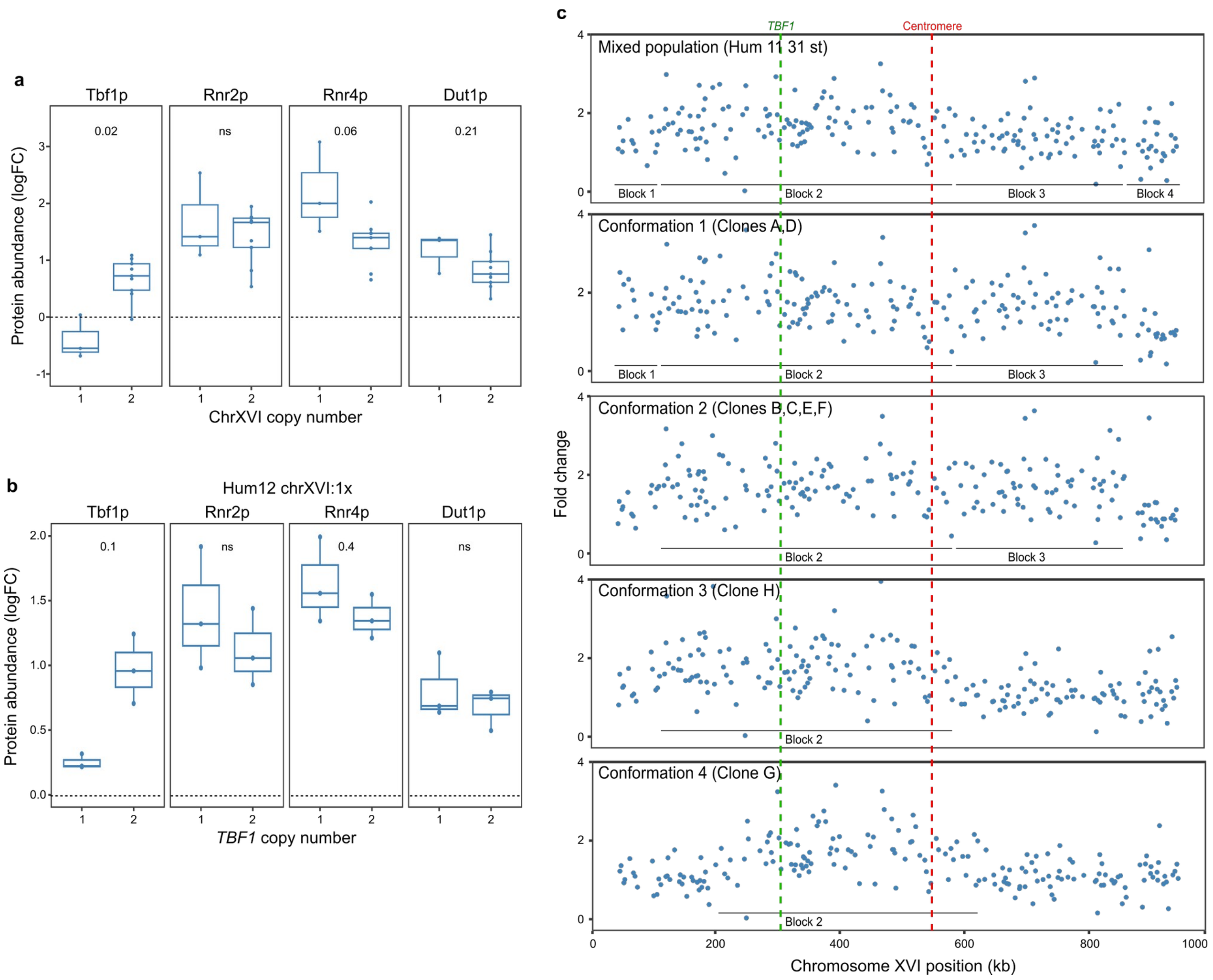

**Supplementary Figure 8 – Protein abundance dynamics upon chromosome XVI aneuploidy and *TBF1* duplication. a-b**, Protein abundance ($log_2FC$) of Tbf1p, Rnr2p, Rnr4p and Dut1p in humanized MAL at 100 scb, carrying 1 ($n$=3) or 2 ($n$=9) copies of chromosome XVI (**a**), and in a humanized line at 100 scb, carrying 1 or 2 *TBF1* copies but only one chromosome XVI (**b**). Boxplots as in Fig. 2. In **a**, humanized lines that have lost the chrXVI aneuploidy before the proteomics measurement ($n$=3) have been excluded from this analysis. Dashed lines indicate the protein abundance (logFC) of the reference strain (wild type at 2 scb, MJD5). Protein abundance measurements were performed in triplicate. **c**, Fold Change profiles (normalized using the wild type at 2 scb) of chromosome XVI for a AE line started from 100 scb (top panel) and of 8 single clones derived from it (bottom panels). FC patterns mirror the conformations shown in Figure 5d. Dashed lines indicate the position of *TBF1* (green) and the centromere (red). Plain lines indicate the length of duplicated blocks as defined by the points of FC breakage in the mixed population (top panel). Chromosome XVI conformations (1 to 4) are ordered descending based on the proportion of chromosome remaining in duplicated state.

**List of primers, oligos and gRNAs used in this study**

| Name | Sequence (5'-3') | Scope |
|---|---|---|
| MEC1_FW | CCTTTCGAACAATTCAGCGTCC | PCR for Sanger sequencing |
| MEC1_RV | GGTATTCAACAGCCAAACTTAGC | PCR for Sanger sequencing |
| TEL1fwSeq | GCCTTTGGTCCTAATGGAAC | PCR for Sanger sequencing |
| TEL1.2_RV | GGTTCGAGAATGATGACAATAGC | PCR for Sanger sequencing |
| FW int MRE11 | CATTACCATTGAATGCGAAA | PCR for Sanger sequencing |
| RV int MRE11 | CAATAGATTACCAAGTTGAA | PCR for Sanger sequencing |
| FW int RAD50 | CAGTTGCTAATGAAAAAGACTA | PCR for Sanger sequencing |
| RV int RAD50 | CATTATACTGATCTATCTTAG | PCR for Sanger sequencing |
| FW MRE11+FW URA3 | ATTAAGAGAATGCAGACAATTGACGCAAGTTGTACCTGCTCAGATCCGATAAAACTCGACTATGTCGAAAGCTACATATAA | Replacement of MRE11 by URA3 |
| RV MRE11+RV URA3 | TCGCGAAGGCAAGCCCTTGGTTATAAATAGGATATAATATAATATAGGGATCAAGTACAATTAGTTTTGCTGGCCGCATC | Replacement of MRE11 by URA3 |
| control ext MRE11 | CTAACTAAGGAAGAAAATTAC | PCR to check mre11::URA3 |
| URA3_Int1 | TGCACGAAAAGCAAACAAAC | PCR to check mre11::URA3 |
| URA3_Int2 | AATGCGTCTCCCTTGTCATC | PCR to check mre11::URA3 |
| Repair cassette Q447P FW | TTTCTCATCTTTATCTACAAACTTCTTTACTGCTTCATTCAAACCAACTTCTGGTAATAAAGATAGTTGCATTTTGTTCAAGAGATCATTAACTAAAGTTGGAACTTCTAGTTCACCGCC | Replacement cassette for CRISPR/Cas9 genetic engineering of Q447P |
| Repair cassette Q447P RV | GGCGGTGAACTAGAAGTTCCAACTTTAGTTAATGATCTCTTGAACAAAATGCAACTATCTTTATTACCAGAAGTTGGTTTGAATGAAGCAGTAAAGAAGTTTGTAGATAAAGATGAGAAA | Replacement cassette for CRISPR/Cas9 genetic engineering of Q447P |
| MRE11 gRNA | TTTCGAATAAACACACATAAACAAACAAAAGTTCACTGATGAGTCCGTGAGGACGAAACGAGTAAGCTCGTCTGAACTAGAAGTTCAAACTTGTTTTAGAGCTAGAAATAGCAAGTTAAAATAAGGCTAGTCCGTTATCAACTTGAAAAAGTGGCACCGAGTCGGTGCTTTTGGCCGGCATGGTCCCAGCCTCCTCGCTGGCGCCGGCTGGGCAACATGCTTCGGCATGGCGAATGGGACACAGGCCCCTTTTCCTTTGTCGATA | Guide RNA for CRISPR/Cas9 genetic engineering of Q447P |
| FW HO+FW TBF1 | TTCATCCAAAATATTAAATTTTACTTTTA | Replacement of HO by TBF1 |

| Name | Sequence (5'-3') | Scope |
|---|---|---|
| | TTACATACAACTTTTTAAACTAATATAC ACATTCTACATCCCATCTTCTAAAT | |
| RV HO+RV TBF1 | CAACTATTAGCTCTAAATCCATATCCTC ATAAGCAGCAATCAATTCTATCTATACT TTAAAATGGATTCGCAAGTGCCCAA | Replacement of HO by TBF1 |
| p132 HO check FWD | CAAATCAGTGCCGGTAACGCT | PCR to check ho::TBF1 |
| HO_gRNA-1_RB_6RC | TTTCGAATAAACACACATAAACAAACAA AAAAATACTGATGAGTCCGTGAGGACG AAACGAGTAAGCTCGTCTATTTTTATAA AGATTGGAGGTTTTAGAGCTAGAAATAG CAAGTTAAAATAAGGCTAGTCCGTTATC AACTTGAAAAAGTGGCACCGAGTCGGT GCTTTTGGCCGGCATGGTCCCAGCCTCC TCGCTGGCGCCGGCTGGGCAACATGCTT CGGCATGGCGAATGGGACACAGCGCAA ATCTTTACTGAT | Guide RNA for CRISPR/Cas9 genetic engineering of HO locus (first strand) |
| HO_gRNA-2_RB_6RC | GGACACAGCGCAAATCTTTACTGATGAG TCCGTGAGGACGAAACGAGTAAGCTCGT CTAAAGACATCGCAAACGTCAGTTTTAG AGCTAGAAATAGCAAGTTAAAATAAGG CTAGTCCGTTATCAACTTGAAAAAGTGG CACCGAGTCGGTGCTTTTGGCCGGCATG GTCCCAGCCTCCTCGCTGGCGCCGGCTG GGCAACATGCTTCGGCATGGCGAATGGG ACACAGGCCCCTTTTCCTTTGTCGATATC AT | Guide RNA for CRISPR/Cas9 genetic engineering of HO locus (second strand) |
| qPCR FW TBF1 (1pair) | CCTTCTCCGATGGAGTCAAA | PCR to check ho::TBF1<br>qPCR chrXVI block 2 |
| qPCR RV TBF1 (1pair) | AGAGTTTCAGGACGAGACG | PCR to check ho::TBF1<br>qPCR chrXVI block 2 |
| FW TBF1+FW cassette | CATCTGCGACAGAAATCATTTTACATTA GTTTTTCTTTACTCACTTCTTTCCCCATTT GTCGACATGGAGGCCCAGAATAC | Replacement of TBF1 by HYGMX |
| RV TBF1+RV cassette | CAAAGTATTTTTAATAGTGAGTGGTTCA ATAGAAAATTAGAAGTCTGTTCATTTAA GCGCGCAGTATAGCGACCAGCATTC | Replacement of TBF1 by HYGMX |
| TBF1 ext ctrl | GACCAGGTTGCTAGAAGATAGG | PCR to check tbf1::HYGMX |
| H2 | CGGCGGGAGATGCAATAGG | PCR to check tbf1::HYGMX |
| H3 | TCGCCCGCAGAAGCGCGGCC | PCR to check tbf1::HYGMX |
| III_PWP2_RT_FW | GTCCCAAACTTTGATTTTCCC | qPCR chrIII control |

| Name | Sequence (5'-3') | Scope |
|---|---|---|
| III_PWP2_RT_RV | TATATCTTGAAGCAGCAGGGC | qPCR chrIII control |
| FUM1 FW | GTTAGTCACCGCTTTGAACC | qPCR chrXVI block 1 |
| FUM1 RV | TGTGTTCAGGAACAACCCAT | qPCR chrXVI block 1 |
| XVI_RPC40_RT_FW | GAAACCGTATCCTTCCTAGCA | qPCR chrXVI block 3 |
| XVI_RPC40_RT_RV | CAAAGGTGAAAGTGCCAGAA | qPCR chrXVI block 3 |
| NUT2 FW | GCCACTACTCAAGACCAAGT | qPCR chrXVI block 4 |
| NUT2 RV | AGCGATCCACATTTCTTTGC | qPCR chrXVI block 4 |
